# Single-cell transcriptomics reveals colonic immune perturbations during amyloid-β driven Alzheimer’s disease in mice

**DOI:** 10.1101/2024.01.27.573841

**Authors:** Priya Makhijani, Rohini Emani, Carlos Galicia Aguirre, Wei-Chieh Mu, Anand Rane, Jenny Hong Yu Ng, Taylor R. Valentino, Max Manwaring-Mueller, Christopher Ryan Tan, Huixun Du, Fei Wu, Saad Khan, Kenneth A. Wilson, Shawn Winer, Chao Wang, Arthur Mortha, David Furman, Lisa M. Ellerby, Olga L. Rojas, Julie K. Andersen, Daniel A. Winer

## Abstract

The “gut-brain axis” is emerging as an important target in Alzheimer’s disease (AD). However, immunological mechanisms underlying this axis remain poorly understood. Using single-cell RNA sequencing of the colon immune compartment in the 5XFAD amyloid-β (Aβ) mouse model, we uncovered AD-associated changes in ribosomal activity, oxidative stress, and BCR/plasma cell activity. Strikingly, levels of colon CXCR4^+^ antibody secreting cells (ASCs) were significantly reduced. This corresponded with accumulating CXCR4^+^ B cells and gut-specific IgA^+^ cells in the brain and dura mater, respectively. Consistently, a chemokine ligand for CXCR4, CXCL12, was expressed at higher levels in 5XFAD glial cells and in in silico analyzed human brain studies, supporting altered neuroimmune trafficking. An inulin prebiotic fiber diet attenuated AD markers including Aβ plaques and overall frailty. These changes corresponded to an expansion of gut IgA^+^ cells and rescued peripheral T_regs_ levels. Our study points to a key glia-gut axis and potential targets against AD.

**Study Highlights:**

- AD is associated with altered immune parameters in the gut of *5XFAD* mice.
- 5*XFAD* colon has reduced ASCs, including CXCR4^+^ cells with a migratory gene signature.
- *5XFAD* brain gliosis includes increased CXCL12 expression.
- CXCR4^+^ B cells and gut-specific IgA^+^ ASCs accumulate in the *5XFAD* brain and/or dura mater.
- Inulin diet attenuates AD disease parameters while boosting IgA^+^ cell and T_reg_ levels.

## Introduction

Late-onset Alzheimer’s disease (LOAD or AD) primarily occurs in individuals over the age of 65, presents with symptoms of dementia and anxiety, and is more prevalent in females [1]. The neuropathology of AD involves an early pre-symptomatic period that correlates with extracellular deposition of amyloid-β (Aβ) followed by a clinical stage marked by the addition of intraneuronal aggregations of hyperphosphorylated tau protein [2,3]. The local innate immune response to Aβ drives microglia activation [4] that can be protective at early stages [5,6]. The early adaptive immune response is also considered to be protective in AD [7]. However, chronic activation of certain microglial subtypes promotes neurodegeneration via orchestrating synapse loss [8], blood-brain-barrier (BBB) breakdown [9,10], tau seeding [11], and infiltration of cytotoxic T cells [12].

The Aβ-associated preclinical phase of AD can persist up to 20 years [13]. This stage has been the subject of intense study to better understand the underlying risk factors, identify early diagnosis biomarkers, and to implement preventative interventions. Interestingly, this phase also correlates with metabolic and gastrointestinal complications. Patient studies have shown heightened prodromal weight loss [14], alterations in energy metabolism [15,16], intestinal dysfunction [17] and changes in gut microbial composition [18,19], compared to healthy aging individuals. An AD-associated loss of intestinal barrier integrity has also been suspected due to elevated levels of fecal proteins in patient serum [20]. A relationship between gut health and neuroinflammation has been noted in several neuropsychiatric conditions and after brain injury [21,22].

Termed the “gut-brain axis”, bidirectional gut-brain interactions drive both protective and deleterious effects in the brain in mouse models of various diseases. For instance, gut-microbiota derived metabolites accumulating in the blood attenuate microglia activation via their ability to permeate the BBB [23–25]. Pre-existing gut dysbiosis promotes neuroinflammation via migratory activation of adaptive immune cells from the gut to the brain during ischemia [26]. In multiple sclerosis (MS), an autoimmune condition targeting myelin antigens, gut IgA^+^ plasma cell migration through the central nervous system attenuates neuroinflammation via IL-10 production [27]. In fungal infections, gut-derived IgA^+^ cells accumulate in the meningeal border region and defend against pathogenicity [28]. AD-specific studies using germ-free mice show that microglia- and astrocyte-mediated neuroinflammation is connected to gut microbiome composition [29,30]. A direct translocation of gut microbes in the AD brain via the vagus nerve has been proposed [31] due to the reported loss of gut barrier integrity [32]. In cases of autoimmunity and cancer, bacterial translocation to disease sites has also been reported [33,34].

However, the nature of the gut immune response in AD, especially with respect to the abundant B lineage cells that partake in microbial homeostasis, is not fully understood. Here, we examined the overall landscape of colonic immune cells in AD using high-dimensional approaches. We found evidence of colon immune activation across both the cellular and humoral arms of adaptive immunity. Remarkably, levels of gut CXCR4^hi^ antibody secreting cells (ASCs) were reduced in Aβ-AD modeled mice. Concurrently, we identified a chemokine axis where microglia and astrocytes in the AD brain produce increased levels of the CXCR4 cognate chemokine CXCL12, correlating with an increase in brain-associated B lineage cells expressing CXCR4. Additionally, dietary intervention with inulin fiber, known to produce microbial-derived short-chain fatty acids (SCFAs) and fortify the intestinal barrier [35,36], attenuated brain Aβ accumulation, gliosis-associated CXCL12 production and CD45^+^ infiltration while boosting T_regs_ and IgA^+^ cell levels in the gut and spleen. Overall, our findings uncovered key alterations to gut and brain immune cells in AD, as well as a likely chemokine axis shared by these cells. Results from our studies also suggest that targeting gut inflammatory responses via dietary and therapeutic interventions could promote both brain and gut health in AD.

## Materials and Methods

### Mice

*C57BL*/6J (000664) and *5XFAD* hemizygous (008730) (B6.Cg-Tg(APPSwFILon, PSEN1*M146L*L28V)6799Vas/Mmjax, MMRRC strain #034848) mice were purchased from The Jackson Laboratory. *5XFAD*-derived *WT* littermates were generated via crossing of hemizygous mice and were housed together after weaning and aged until specified harvest dates. Mice were maintained under pathogen-free, temperature and 12h light-dark cycle-controlled environments at the Buck Institute for Research in Aging, Toronto Medical Discovery Tower, or Krembil Brain Institute’s vivarium facilities. The animal facility at the Buck Institute for Research on Aging is accredited by AAALAC International (Unit Number 001070). All protocols and procedures described herein were approved by the Buck’s Institutional Animal Use Committee. Female mice were used for all experiments, unless specified.

### Immune cell isolation

Mice were euthanized using CO_2_ inhalation prior to collecting colons, blood, and spleens. After euthanization, blood was collected via cardiac puncture into K2-EDTA tubes (BD, 367855). Spleens were filtered through 70 μm cell strainers to generate single cell suspensions. Blood and spleens were subjected to hemolysis. Colons were processed using a lamina propria dissociation kit (Miltenyi Biotec, 130-097-410) with the gentleMACS Dissociator (Miltenyi Biotec, 130-093-235). Prior to brain collection, mice were additionally transcardially perfused with 10-20 mL of cold PBS. Mouse whole brains were processed using a brain dissociation kit (Miltenyi Biotec, 130-107-677) with the gentleMACS Octo Dissociator (Miltenyi Biotec,130-096-427). All cell suspensions were filtered through a 40 μm cell strainer prior to downstream application.

### Single-cell RNA sequencing analysis

*<u>Cell sorting and sequencing</u>* Single cell suspensions of colons from 9-month-old littermate *WT* and *5XFAD* female mice were prepared as described above. Cells were stained with LIVE/DEAD Fixable Blue cell stain (Thermofisher, L23105) for viability assessment and blocked with FcBlock with CD16/32 (Biolegend, 101320). Cells were then stained with anti-mouse CD45 (Biolegend, 30-F11). Live CD45^+^ cells were sorted using the FACS Aria sorter at the Buck Institute for Aging Research flow cytometry core facility and collected into RPMI-1640 (Wisent) with 10% FBS (Wisent). Samples were processed utilizing a Chromium Next GEM Single Cell 3’ GEM, Library & Gel Bead Kit v3.1. The resulting libraries were sequenced at UC Davis Genome Center Bioinformatics Core using Illumina NovaSeq 6000 sequencing. The resulting sequencing data was processed using Cell Ranger (6.0.1). *<u>Data Processing.</u>* The Seurat (v4.0.2) workflow was used to filter, down sample, normalize, scale, integrate and cluster (res 0.9), and plot the data. Differential expression was performed utilizing the MAST method in the Seurat function FindMarkers. The SCTransform method was used to identify top differentially expressed genes (DEG) between genotypes. *<u>Trajectory Analysis</u>*. To confirm the putative B cell developmental pathways in the colon, cell trajectory analysis was conducted using the R package, *Monocle 3* (v1.3.4) [37]. The standard *Monocle 3* pipeline with default settings was used without pruning for learn_graph( ). Finally, order_cells( ) function was run using Naive B cells (Cluster 0) as the root of the trajectory [37].

### Flow cytometry

Single cell suspensions of mouse tissue and blood were stained using the LIVE/DEAD Fixable Blue cell stain (Thermofisher, L23105) for viability assessment and blocked with CD16/32 based FcBlock (Biolegend, 101320) for 20 min at 4°C. Cells were further stained with fluorophore-conjugated antibodies for 30 min at 4°C in the dark. The following antibodies were used CD11b-BUV563 (BD Biosciences, clone: ICRF44), CD11b-BV650 (Biolegend, Clone: M1/70), EpCAM-BV711 (Biolegend, Clone: G8.8), CD19-APCCy7 (Biolegend, Clone: 6D5), IgD-BUV605 (Biolegend, Clone: 11-26c-2a), IgM-PeCy7 (Biolegend, Clone: Rmm-1), CD11c-efluor450 (Invitrogen, Clone: N41B), CD45.2-AF700 (Biolegend, Clone: 104), CD127-BV711 (Biolegend, Clone: A7R34), B220-PEFire810 (Biolegend, Clone: RA3-6B2), CD45-BUV805 (BD Bioscience, Clone: 30-f11), CD23-APCCy7 (Biolegend, Clone: B3B4), CD138-BV650 (Biolegend, Clone: 281-2), CD31-APCCy7 (Biolegend, Clone: MEC13.3), LPAM-1(Integrin α4β7)-PE (Biolegend, Clone: DATK32), CXCR4-BV421 (Biolegend, Clone: L276F12), CD86-AF700 (Biolegend, Clone:GL-1), CD86-PeDazzle594 (Biolegend, Clone:GL-1), IgM BUV615 (BD, Clone: R6-60.2), CD21-BV421 (Bioleged, Clone: 7E9), MHCII(I-A/I-E)-BUV395 (BD, 2G9), CD43-PeCy5 (Biolegend, Clone: 1B11), CD11c-BUV737 (BD, Clone: HL3), KLRG1-BV480 (BD, Clone: 2F1), GL7-PerCpCy5.5 (Biolegend, Clone: GL7), gp38-PerCpCy5.5 (Biolegend, Clone: 8.1.1), CD3-APCFire810 (Biolegend, Clone: 17A2), IgM-BV510 (BD, Clone: II/41), CD45-BUV737 (BD, Clone: 30-f11), F4/80-BV480 (BD, Clone: T45-2342), CD19-BV570 (Biolegend, Clone: 6D5). This was followed by fixation and permeabilization using either the FOXP3 staining buffer set (eBioscience, 00-5523-00) or the Cytofix/Cytoperm kit (BD, 554714). Intracellular staining was done using the following antibodies: BLIMP1-PE (Invitrogen, Clone: 5E7), CXCL12-PE (R&D Biosystems, Clone: IC350P), IgA-BV786 (BD, Clone: C10-1), c-JUN-AF647 (Cell Signalling Technologies, Clone: 60A8), CXCR4-BUV737 (BD, Clone; 2B11), FOXP3-AF488 (Biolegend, Clone: 150D), RORγt-BV421 (BD, Clone: Q31-378), RORγt-PercPefluor710 (Invitrogen, Clone: AFKJS-9). Stained cells were analyzed on the Cytek Aurora at the Buck Institute’s Flow Cytometry Core facility within 24h. All raw data was unmixed using SpectroFlo (v3.0) and analyzed using FlowJo (v10.8.1).

### Frailty scoring

The health span of mice was assessed using a 31-term frailty index to identify humane interventions and endpoints. Non-invasive clinical assessment of 31 physical manifestations of age-related deficiencies and scored based on severity was conducted using previously published recommendations [38]. Scoring was performed on mice between 10 and 18 months of age.

### Immunohistochemistry and immunofluorescence microscopy

Colon Swiss rolls and perfused whole brains were fixed with 10% buffered formalin for 24-48 h and embedded into paraffin blocks. FFPE tissues were sectioned into 5 μm slices and mounted onto slides. Following xylene-based deparaffinization, heat-induced antigen retrieval was performed at pH 6.0 (Biocare Medical, Rodent Decloaker) prior to hematoxylin and eosin (H&E) or immunofluorescence staining. The following primary antibodies were used: rabbit anti-mouse IgA (NSJ Bioreagents R20169), mouse anti-mouse CXCL12 (R&D, MAB350), rat anti-mouse CD45 (eBioscience, 14-0451-82), rat anti-mouse CD19 (Invitrogen 13-0194-82), mouse anti-mouse amyloid beta (Clone 6E10, Biolegend, 803004), and rabbit anti-mouse Iba1 (Fujifilm, 019-19741). Appropriate secondaries were used in AlexaFluor 488, 647 or 555 as specified. The following conjugated antibodies were used: rabbit anti-mouse c-JUN AF647 (Cell Signaling Technology, 40502) and rat anti-mouse B220-PE (Biolegend, 103207). Mouse-on-mouse blocking reagents (Vector Laboratories) were used where required. After DAPI (Sigma) staining and mounting (ProLong Gold Antifade Moutant, ThermoFisher, P36930), the Zeiss Axioscan 7 microscope slide scanner was used to collect 20X images. Images were processed using Zen Blue (Zeiss) software for background normalization and Fiji [39] for quantification.

Spleens were placed in a histology tray, embedded in OCT (Fisher Healthcare), and frozen in 2-methylbutane cooled on dry ice. Trays were wrapped in aluminum foil and stored at −80C until further use. Frozen tissues were subsequently sectioned with a cryostat to 5-6 μm in preparation for acetone fixation and staining. Defrosted and PBS-re-hydrated tissues were subjected to FcReceptor blocking (BD Pharmingen, 553141) and primary antibody staining with rat-anti-mouse antibodies. Data depicted were obtained by staining with B220-AF647 (Biolegend, 103226) and CD4-PE (Biolegend, 100512). After DAPI (Sigma) staining and mounting (ProLong Diamond, ThermoFisher, P36965), spleen slides were visualized by microscopy using the Zeiss AxioObserver with Zen Blue (Zeiss) software. Images were processed using Fiji [39].

### Behavioral assays

*<u>Open Field Test</u>.* Mice were placed individually into activity cages equipped with rows of infrared photocells interfaced with a computer (San Diego Instruments). After a 15-min adaptation period, open field activity was recorded for 20 min. Recorded beam breaks were used to measure total path lengths in the margin versus the center of the cage, along with active times and rearing events. *<u>Small y-maze</u>.* Mice were placed in the center of a small Y-maze (arm length: 15 cm), and spontaneous alternation was recorded in a single continuous 6 min trial by a live observer. Each of the three arms was designated a letter A–C, and entries into the arms were recorded. The percent of spontaneous alternation was calculated over the total number of entries possible.

### Dietary Interventions

Fiber diet studies were conducted on age-matched female *WT* and *5XFAD* mice fed *ad libitum* from 8 weeks to 15 of age months with the following diets. Mice were fed cellulose diet (D13081109i, Research Diets Inc.) containing 16.6% cellulose and no inulin; or inulin diet (D13081108i, Research Diets Inc.) containing 17.3% inulin and no cellulose; or a control diet (D12450Ji, Research diet) containing 4.7% cellulose comparable to standard chow.

### Commensal enzyme-linked immunosorbent spot assay

Dura mater was scored from the skull and dissociated with Collagenase P and DNase I in 2% FBS for 20 min at 37° C. After neutralization with 2 mL PBS, cells were counted to perform the assay. The ELISPOT assay was conducted as previously described [27]. Briefly, membrane plates (MiliporeSigma, MSIPS4W10) were coated with 100 μl/well of allogenous heat-killed fecal matter (1mg/mL) diluted 1:10 and placed at 4°C overnight. Plates were then blocked with 100 μl/well of 10% FBS/RPMI (Sigma) for 2 h at 37°C. Starting with 1×10^6^ cells, 200 μl/well of single-cell suspension solutions in FBS/RPMI were loaded onto the plate in serial 2-fold dilutions and left overnight at 37°C. Cells were discarded the next morning, and plates were washed with 0.1% Tween-20 in PBS. Subsequently, HRP-conjugated IgA and AP-conjugated IgG detection antibodies were added and incubated for 2 h at 37°C. Plates were developed using AEC Peroxidase (for HRP-conjugated Ab) (Vector Laboratories, cat. #SK-4200) or Vector Blue (for AP-conjugated Ab) (Vector Laboratories, cat. #SK-5300) substrates to detect IgA and IgG ASCs, respectively. After drying, spots were counted using a light microscope and quantified with respect to the original cell concentration.

### Statistical analyses

Statistical difference between two groups was determined using a two-tailed Welch’s t-test (i.e. assuming unequal standard deviations), with GraphPad Prism Software (v9.4.0). All data was presented as mean +/- SEM. Statistical significance was set at p=0.05, with * denoting p≤0.05, ** p ≤0.01, and *** p≤0.001.

## Results

### Altered intestinal immune composition across the colonic immune landscape in Aβ-driven AD

To identify immunological changes to the gut-brain axis in Aβ-AD, we used the well-studied *5XFAD* model. These mice co-express five familial human AD mutations, all driven by the *Thy1* promoter and resulting in extracellular Aβ deposition and plaque formation in the brain starting at ∼2 months of age and loss of pyramidal neurons by 9 months [40]. Neurodegenerative changes typically exhibit as both cognitive and motor impairments that recapitulate aspects of human disease [41,42]. We observed similar phenotypes to those previously reported in this mouse model [43]. Between 9 and 13 months of age, frailty increased in *5XFAD* compared to *wildtype* (*WT)* age-matched mice (Fig. 1A) on a 31-point frailty index [38]. Frailty changes included a significant reduction in total body weight (Fig. 1B), evident body tremors and instances of diarrhea/constipation (Fig. S1A). This correlated with reduced spleen weight and colon length (Fig. S1B). Increasing anxiety was also observed via the open field behavioral assay in *5XFAD* mice by 12 months (Fig. S1C), though no changes were seen using the Y-maze spontaneous alternations test (Fig. S1D).

**Figure 1:**
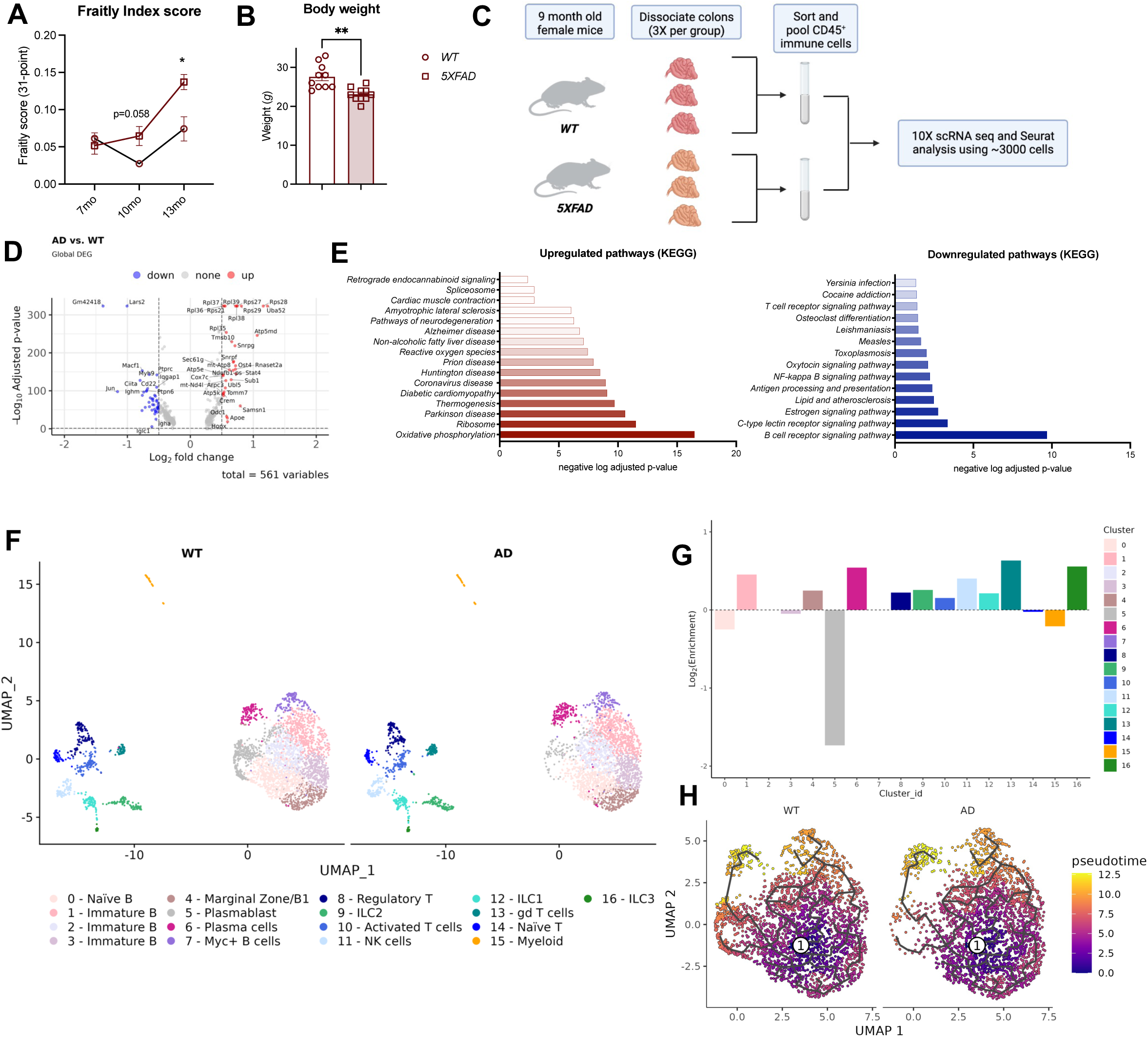
Single-cell sequencing reveals altered immune parameters in Alzheimer’s disease colons. **(A)** 31-point frailty scores comparing *5xFAD* (square) to *WT* (circle) female mice at specified ages, n=5 per group. **(B)** Body weight of 12-month-old *5xFAD* and *WT* mice, n = 10 per group. **(C)** Schematic workflow of scRNA sequencing experiments. **(D)** Volcano plot of differentially expressed genes (DEGs) across all immune cell types between *WT* and *5xFAD*. DEGs (including log2FC and adj.p) were calculated by the MAST method. Plotted adjusted p-value (adj. p) cutoff is 0.05, and log2FC cutoff is 0.5. **(E)** KEGG pathway scores of significantly (adj. p cutoff is 0.05) downregulated (left) and upregulated (right) DEGs evaluated using g:Profiler using an ordered query. **(F)** UMAP plot of colon-derived CD45 immune cell single-cell transcriptomes (10X Genomics) pooled together as described in (C). Cells were resolved into 17 distinct clusters. **(G)** Quantification of relative abundance of each cluster of single immune cells by log2Fold change (FC) between *WT* and *5xFAD* mice. **(H)** UMAP of pseudotime trajectory analysis of *WT* and *5xFAD* colon B cells. White circles represent the root node designated at the germinal center (cluster 7). Black lines depict branch pathways, where cells travel to a variety of predicted outcomes from intersecting points.

In this study, we focused on the colon because it has higher levels of bacterial colonization than the small intestine [44], and bacteria-derived metabolites are thought to affect neuroinflammation and vice versa [45]. Using H&E staining, *5XFAD* mice showed no overt signs of acute or chronic colonic inflammation (Fig. S1E). Colon weight-to-length ratios, used in colitis to measure overt gut inflammation [46], also showed no significant differences between groups (Fig. S1B). Changes in spleen weight were correlated with body weight decreases and we observed no changes in splenic follicle number or T and B cell area in the *5XFAD* mouse (Fig. S1F). By immunofluorescence (IF) staining, we found no Aβ in the *5XFAD* colon (Fig. S2) as previously reported [47].

For a deeper investigation of AD-associated compositional immune perturbations in the colon, we performed single-cell RNA sequencing (scRNAseq) analysis on sorted CD45^+^ immune cells from three collagenase-dissociated colons of 9-month-old female *5XFAD* and three age-matched littermate *WT* mice (Schematic Fig. 1C). We used 9-month-old *5XFAD* mice to study stable gut immune changes at a time-point when neurodegenerative outcomes are well-established. Since more AD patients are female [48], and females more strongly manifest disease in the *5XFAD* mouse model [49], we focused our study on female mice. Using these experimental conditions, we created a single-cell map of the colon immune landscape that comprised of 3443 cells from *5XFAD* and littermate *WT* control mice.

Across all immune populations in the colon, *5XFAD* altered expression of 561 genes out of which 550 genes had a significant adj. P-value cutoff of <0.05. The total list of significant differentially expressed genes (DEGs) is visualized using a volcano plot (Fig. 1D), highlighting the most highly changed transcripts (|logFC| > 0.5). We observed pan-immune induction of ribosomal genes (*Rps*/40S family, *Rpl*/60S family, *Eif* family), mitochondrial genes (*Cox7c*, *Tomm7*) and immune activation genes (*Lyn*, *Stat4, H2*/histocompatibility family) in *5XFAD* colon immune cells (Supplementary Data 1). Globally downregulated genes included *Igha*, *Ighm*, *Ighd*, *Cd79b*, *Cd22*, *Irf8*, and *Ciita* normally associated with B cells (Fig. 1D). The DEGs with adj. P-values <0.05, separated into up and downregulated genes by logFC value were entered into g:Profiler as an ordered query (Supplementary Data 2). Resulting KEGG pathways upregulated in the *5XFAD* mice included neurodegenerative pathways (amyotrophic lateral sclerosis, Huntington’s, Parkinson’s, prion and Alzheimer’s disease), and innate immune inflammation (ribosome, reactive oxygen species, oxidative phosphorylation, COVID-19) (Fig. 1E, left). KEGG pathways downregulated in the *5XFAD* mice included adaptive immune-associated genes, most significantly BCR as well as TCR and NF-κB signaling (Fig. 1E, right). Overall, perturbations in protein synthesis, metabolism, and immune activation reflect the manifestation of AD-associated pathology in gut immune cells.

To take a closer look at immune cell subtype-specific changes, we used dimensionality reduction to identify 17 clusters of immune cells visualized by Uniform Manifold Approximation and Projection (UMAP) (Fig. 1F). Based on the presence of canonical markers (Fig. S3A-B) we identified one myeloid, eight B lineage, and eight T cell/innate lymphoid cell (ILC) clusters (Fig. 1F). DEGs in individual clusters are shown in Fig. S3C and Supplementary Data 3. Minimal to no changes in proportions of T lineage clusters were observed in *5XFAD* colons and innate cell clusters, including myeloid and ILC lineages, at the assessed resolution (Fig. 1F-G). Most remarkably, we observed a near complete reduction in the frequency of B lineage cluster 5 in the *5XFAD* colons, with lesser changes in other B clusters (Fig. 1F-G). We identified cluster 5 as a mixed cluster representing plasmablasts (PBs) and maturing plasma cells (PCs) due to distinguishing transcriptional features lacking in the other seven B lineage clusters. The partial lack of *Vpreb3* (surrogate light chain) and a presence of lambda light chain (*Iglc2*) transcript suggest that these cluster 5 cells have likely completed VDJ recombination [50] (Fig. S3B). Further, the expression of *Cxcr4*, *Fos*, *Fosb*, *Jun*, *Irf4*, *Irf8*, *Blc11a* and partially *Spib*, *CD38*, *CD22*, *Ciita*, *Ms4a1*, *Cd79b*, *Pax5*, *Ighm* and *Ighd* suggests that cluster 5 is differentiating through developmental checkpoints towards a mature antibody-secreting plasma cell [51] (Fig. S3B). A fully mature PC population appeared in nearby cluster 6 with *Aicda* transcript, responsible for class switch recombination, and *Prdm1*, *Zbtb20*, *Xpb1*, *CD80*, *Igha*, and *Ighg2b* mRNAs [51] (Fig. S3B). B cell trajectory prediction analysis conducted using *Monocle3* showed that pseudotime starting at naïve B cells (cluster 0) results in some cells differentiating into PCs (cluster 6), via several possible paths including a possible germinal center-like Myc^+^ B cell (cluster 7) [52](Fig. 1H, S4A-B). PB cluster 5 cells were predicted to partially give rise to PC cluster 6, with developmental trajectories remaining consistent between *WT* and *5XFAD* colon B cells (Fig. 1H, S4A-B). Although this did not rule out stress-induced apoptosis of any short-lived PBs within cluster 5, an abundance of migratory signatures [53] within this cluster and previous reports of neuroinflammation driven gut-brain PC migration [27,54] support plausible colon PB emigration in the *5XFAD* model.

### Loss of CXCR4^hi^ gut plasma cells and systemic B cell activation in *5XFAD*

The striking loss of cluster 5 B cells in *5XFAD* colons led us to take a closer look at the transcriptional signatures in all PB/PC subsets. Collectively termed antibody secreting cells (ASCs), cluster 5 (PB, or maturing PC) and cluster 6 (mature PC) showed transcription of several migratory receptors thought to respond to a variety of signals, including *Cxcr4, Ackr3* (CXCR7), *Tnfrsf13c* (BAFFR), *Gpr183*, *Cd38*, *S1pr1, Ffar1, Gpr174, and Itgb7* (Fig. 2A, S3B). A similar expression signature was recently reported in *Kaede* mice including *Cxcr4, Itgb7, Gpr174* and, transiently AP-1 factor, *Jun,* which defined a gut-imprinted transcriptomic signature in colon B cell emigrants in the spleen under homeostatic conditions [53]. Isolated to the receding cluster 5, we also found *Klf* family genes (*Klf2, Klf4, Klf5, Klf6*) (Fig. 2A, Fig. S3B), transcription factors that regulate B cell migratory programs [55] and IgA responses [56]. Under pathological conditions, gut-specific ASCs emigrate to sites of peripheral inflammation [57,58], especially the CNS [27,59]. As such, the loss of cluster 5, displaying a migratory signature, points towards possible colon ASC emigration in AD.

**Figure. 2:**
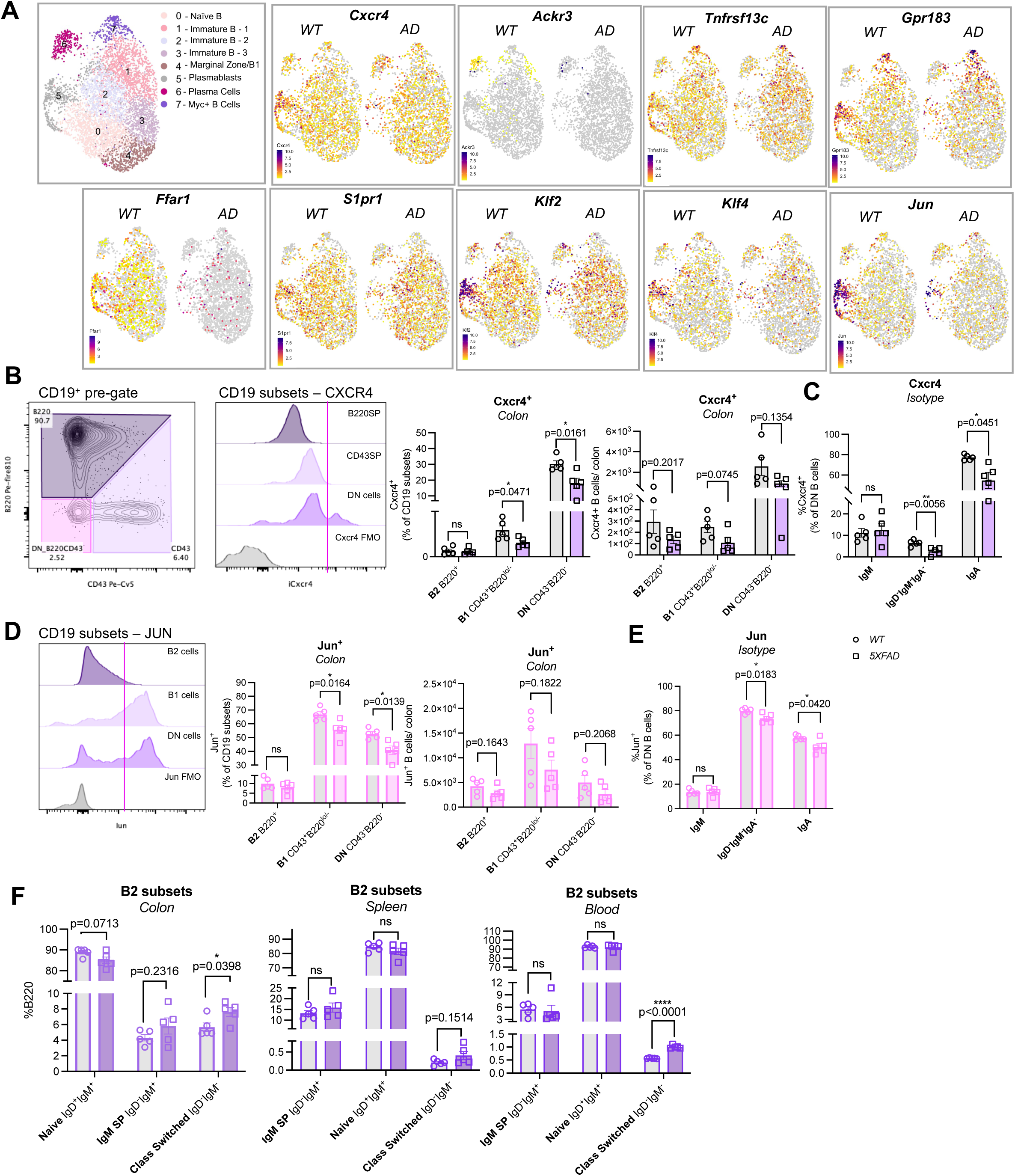
Alzheimer’s disease changes the B lineage landscape within the colon. **(A)** UMAP showing transcription changes in key migratory genes in *WT* and *5xFAD* colon B cells**. (B)** Flow cytometric classification of CD45^+^CD19^+^ B cell subsets in 12mo female mouse colons (left). Representative histograms of CXCR4 expression (gMFI) per subset in *WT* and *5xFAD* (middle) quantified by both gated frequency and cell number (right) in *WT* (circle) and *5xFAD* (square) mice; n = 5 per group. **(C)** Frequency of CXCR4^+^ cells per isotype within the B220^-^CD43^-^ (DN) population. **(D)** Representative histograms of JUN expression (gMFI) per subset in *WT* and *5xFAD* (left) quantified by both gated frequency and cell number (right); n = 5 per group. **(E)** Frequency of JUN^+^ cells per isotype within the B220^-^CD43^-^ population. **(F)** Frequency of naïve, IgM and class switched B2 cells within the B220^+^CD43^-^ SP (B2 cells) population in colon, spleen and blood.

We next corroborated these transcriptional signatures with protein expression in colon B lineage cells using high-dimensional spectral tuning flow cytometry. First, we classified broad B cell subsets using CD43 and B220 expression within the CD19 immune compartment to capture B lineage heterogeneity within the colon (gating strategy; Fig. S5). We used CD19^+^CD43^+^B220^lo/-^ (CD43 single positive; SP) to mark B1 cells, though activated B and PCs may also exist in this population [60] (Fig. 2B). We used CD19^+^CD43^-^B220^+^ (B220SP) to represent the majority of B2 cells (Fig. 2B). Finally, because the traditional PC marker CD138 is sensitive to colon dissociation protocols, we used the CD43^-^B220^-^(double negative; DN) population to locate CD19^+^ maturing ASCs (Fig. 2B). We suspect that, because the CD19-BCR complex is no longer required in mature PCs, the CD19^+^B220^-^CD43^-^ DN gate captures mainly early plasma cells but cannot rule out transitioning plasmablasts. We used the term ASC to represent these multiple possibilities. Indeed, upon sub-gating, we found that DN cells were class switched (IgD^-^IgM^-^) or IgM^+^SP, with most of the class switched cells being either IgA^+^ or IgM^-^ IgA^-^, the latter likely representing IgG isotypes (Fig. S5). Consistently, B220SP cells were largely naïve (IgM^+^IgD^+^), or class switched (Fig. S5). To measure mature PCs, we used terminal marker BLIMP1 [61] in the CD3^-^CD19^-^ populations and found a significant reduction in the frequency of BLIMP1^+^ cells in the *5XFAD* colon (Fig. S6A). Also, total gut IgA^+^ cells showed a trending decrease in *5XFAD* by flow cytometry (Fig. S6B) and immunofluorescence staining (Fig. S6C), consistent with single cell RNA expression analysis.

We then investigated CXCR4 protein expression in each B lineage subset, as per our scRNAseq findings. The proportions of CXCR4^hi^ cells were significantly reduced within the CD43SP and DN compartments, with trending decreases in cell number in *5XFAD* compared to *WT* colons (Fig. 2B). Interestingly, the DN cells had the highest proportion and levels of CXCR4 expression compared to B220 SP and CD43 SP populations (Fig. 2B). Within this DN group, the IgA isotype had the highest expression of CXCR4 compared to others (Fig. 2C), suggesting a stronger ability to respond to CXCR4-mediated chemotaxis than other B lineage cells. Consistently, we found a significant decrease in frequencies of CXCR4^+^ ASCs both within the IgA^+^ and IgM^-^IgA^-^IgD^-^ isotypes (Fig. 2C) together with a trending decrease in cell number in *5XFAD* colons (Fig. S6D). Other studies have shown that colon ASCs are largely IgA^+^ while expressing high levels of CXCR4^+^ and IL10 [62]. We also checked for the expression of JUN, found in developing plasmablasts [63,64] and early CXCR4 signaling [65,66]. We found reduced JUN levels within both CD43SP and DN subsets (Fig. 2D), as well as within the IgA^+^ and IgM^-^ IgA^-^IgD^-^ isotype subsets (Fig. 2E, Fig. S6E) in *5XFAD* compared to *WT* colons. The location of CD19^+^JUN^+^ B cells [67] in the colon was validated by immunofluorescence staining; they mapped to the outskirts of isolated lymphoid follicles (ILFs) (Fig. S6F). Overall frequencies and cellularity of total B cells and of each B cell subset (CD43SP, B220SP, DN) were unchanged in AD colons by flow cytometry (Fig. S6G-H), likely reflecting the reported rapid replenishment of colon B emigrants to maintain gut immune homeostasis [53].

As predicted, we also observed an increase in class switched B220SP cells and a trending decrease in naïve B220SP cells in *5XFAD* colons (Fig. 2F). By scRNAseq, we observed activation signatures across all colon B cell clusters (0-7) (Fig. S7). Transcriptionally upregulated genes included *Stat4*, *Cd86, H2-Ab1, H2-D1, H2-Eb1, Lyn, Cd86* associated with immune activation, and *Eif4a1*, *Rps28*, *Rpl38*, *Uba52*, *Cox7c*, *Tomm7*, *mt-Atp8* associated with translation by ribosomes and metabolic activity by mitochondria, representing pan B cell dysfunction in the *5XFAD* colon (Fig. S7). By flow cytometry, an increased proportion of class switched B2 cells in AD colons was mirrored in the blood, with trending increases in the small population of class switched B2 cells in the spleen (Fig. 2F, gating strategies Fig. S8A-B). With increased B2 class switching, an increased proportion of DN B cells (CD19^+^B220^-^CD43^-^) was observed in *5XFAD* spleens with a trending increase in the blood compared to controls (Fig. S9A-B), indicative of systemic B cell changes. Consistently, an increasing trend in the frequency of IgA^+^ cells was observed in blood with no differences in the spleen (Fig. S9C). Trending increases in the expression of activation marker CD86 in splenic B2 cells and increased MHCII levels in blood B2 cells were observed in *5XFAD* mice (Fig. S9D-E). We also noted other markers of systemic B cell dysfunction, including increased marginal zone B cell (MZB) proportions in *5XFAD* spleens (Fig. S9F), likely indicating increased antigen encounter.

Taken together, the reduced amounts of migratory gene expressing colon ASCs, increased B2 cell class switching and inflammatory markers throughout the periphery suggest whole-body B cell dysfunction in *5XFAD*, with the possibility of ASC emigration from the colon. The colon B activation observed here may be linked to a dysbiosis phenotype [32,68,69], and the peripheral B cell immunological changes may be correlated with the leaky gut [31] and changing blood BCR repertoire observed in AD [70].

### AD CNS contains CXCL12-producing glial cells, coupled to increased CXCR4^+^ B cells, and IgA^+^ ASCs that recognize gut commensal antigens

Next, we sought to better understand the effect of systemic inflammation, including neuroinflammation, correlating with the observed gut ASC phenotype. There was a transcriptional downregulation of CXCL12 binding receptors; its cognate receptor, *Cxcr4*, and sequestration receptor, *Ackr3* (CXCR7) were found in cluster 5 (PB) and cluster 6 (PC), respectively (Fig. 2A). Although CXCR4 is also expected to bind to MIF and CXCL14, it preferentially responds to CXCL12 chemotaxis [71,72]. Additionally, B cell migration has been linked to CXCR4-mediated trafficking [73] specifically in the brain [74,75] with CXCR4 expression reported in dura mater IgA^+^ cells [28]. Therefore, we predicted that this phenotype exists in AD and is orchestrated by a CXCL12 signal. To study this effect, we conducted high-dimensional spectral tuning flow cytometry analysis on dissociated mouse brains from *5XFAD* mice (gating strategy; Fig. S10). We first confirmed that activated microglia accumulate in the *5XFAD* brain at later stages [76]. We observed an overall increase in the number of microglia in the AD brain marked by CD11b^+^CD11c^+^ within live CD45^med^ cells (Fig. 3A), and an increase in intracellular CXCL12 expressed by those microglia (Fig. 3B). Using *in silico* analysis of published human single-cell dataset [77], we identified increases in CXCL12 transcript in a subset of AD microglia and astrocytes versus healthy aging in prefrontal cortex tissue (Fig. 3C). By immunofluorescence (IF) tissue staining we found that a proportion of *5XFAD* microglia (CD45^lo^) expressed high levels of CXCL12, specifically in the dentate gyrus region compared to *WT*. IF also revealed increased CXCL12 expression on CD45^-^ astrocyte-shaped cells, with enrichment seen in subregions of the hippocampal formation (Fig. 3D). *WT* brains appeared to only produce CXCL12 in rounded structures, morphologically consistent with endothelial cell vasculature (Fig. 3D). Minor or trending increases in total levels of adaptive immune cells (CD45^hi^) were seen in *5XFAD* brains (Fig. S11A-B). By flow cytometry, we also saw a decrease in brain endothelial CD31^+^ cell frequencies and reduced CXCL12 production by endothelial cells in *5XFAD* brains (Fig. S11C-D). Overall, our findings showed a significant increase in gliosis-associated CXCL12 expression in the *5XFAD* brain, with a suspected aberrant expression pattern in endothelial cells, pointing towards dysfunctional trafficking of CXCR4^+^ cells.

**Figure 3:**
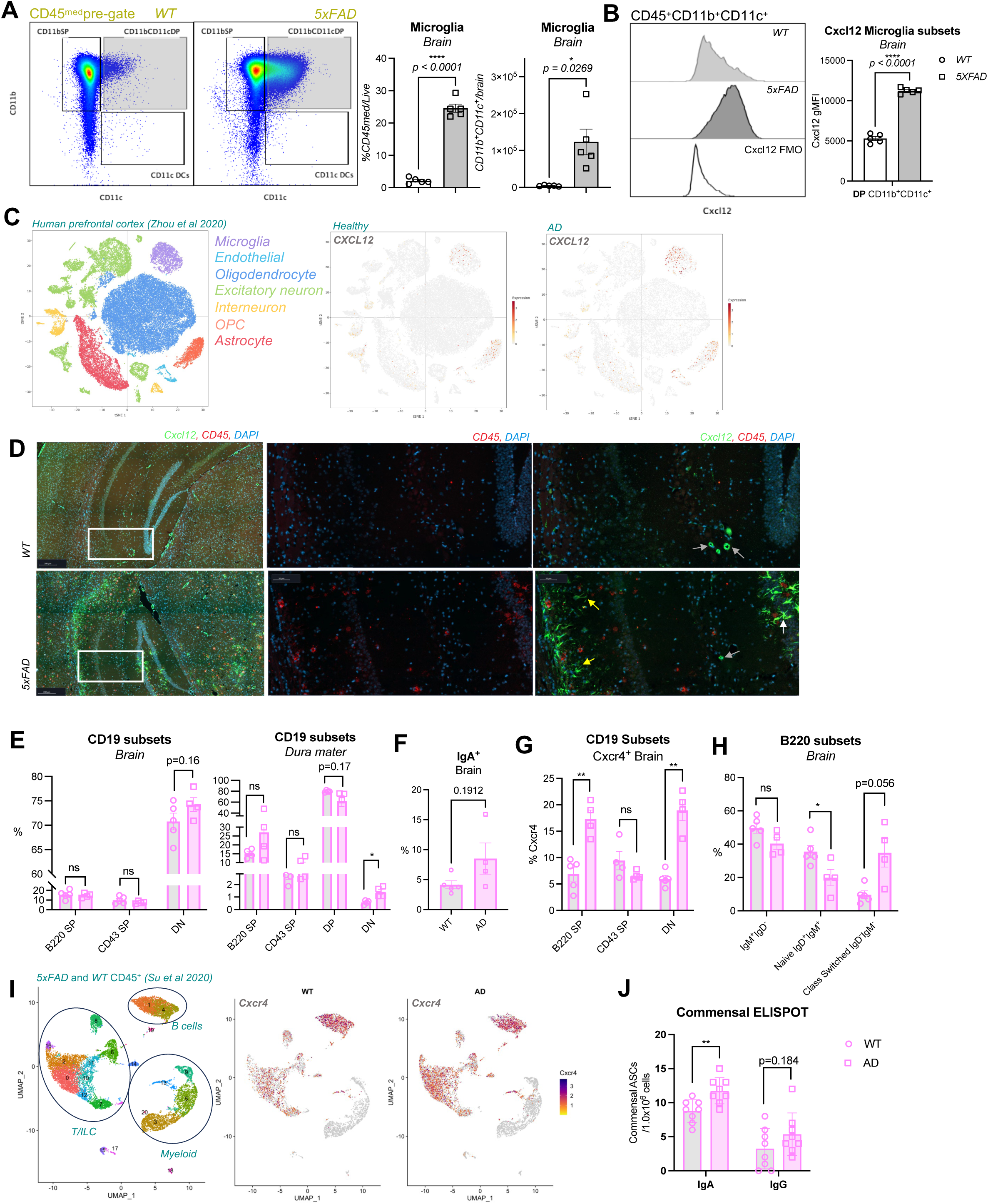
Increased CXCL12 production by AD glial cells correlates with brain and dura mater infiltration of CXCR4^+^ immune cells. **(A)** Representative plots of flow cytometric assessment of myeloid lineage cells in the brain parenchyma (left). Frequencies and total cell numbers of microglia marked using CD11b^+^CD11c^+^ population in *WT* (circle) and *5xFAD* (square) 12-month-old female mice; n = 5 per group. **(B)** Representative histograms (left) and quantification of CXCL12 gMFI within the microglia population (right). **(C)** Representative immunofluorescence staining of brain hippocampal regions with intracellular CD45 (red) and CXCL12 (green) antibodies in *WT* (top) and *5xFAD* (bottom) 12mo female mice. CXCL12 expression in endothelial structures (grey arrow), CD45^-^ astrocytic cells (yellow arrow), and CD45^+^ microglia (grey arrow) are highlighted. **(D)** UMAP of reanalyzed human post-mortem prefrontal cortex CD45 scRNAseq analysis from Zhou *et al.* 2020 (left) with *Cxcl12* transcription in healthy and AD tissue (right). **(E)** Frequencies of CD19^+^ B cell subsets evaluated by B220 and CD43 expression using flow cytometry analysis of brain and dura mater cells from *WT* (circle) and *5xFAD* (square) 12mo female mice; n = 5 per group. **(F)** Total IgA^+^ cells within the brain parenchyma of *WT* (circle) and *5xFAD* (square) 12mo female mice measured using flow cytometry; n = 5 per group. **(G)** Commensal ELISPOT assessment number of IgA and IgG ASCs in the dura mater of 9mo female mice; n = 8 per group. **(H)** Flow cytometric assessment of frequencies of Cxcr4^+^ B cells in the brain using CD19^+^ subsets (left) and CD19^+^B220^+^CD43^-^ SP class switched B2 subsets (right). **(I)** UMAP of reanalyzed mouse brain CD45 scRNAseq analysis from Su *et al.* 2023 (left) with *Cxcr4* transcription in 8mo female *WT* and *5xFAD* mice (right).

Next, we sought to investigate B cells and ASCs within the brain and brain border regions. We found increases in B220^-^CD43^-^ (DN) CD19^+^ B cells within the AD brain and dura mater (Fig. 3E), as well as a trending increase in frequency of IgA^+^ ASCs expected to be of mucosal origin (Fig. 3F). Within the AD brain parenchyma, flow cytometry analysis showed increases in CXCR4^+^ B cells, including in both B220SP and DN subsets (Fig. 3G). We also observed a near significant increase in class switched B220SP B2 cells in the brain, along with a decrease in naïve B cells (Fig. 3H). Data mining of *5XFAD* brain immune cell scRNAseq data from Su et *al*. 2023 [7] and Keren-Shaul et *al.* 2017 [4] showed that brain-infiltrating B and T lineage cells also express *Cxcr4* mRNA at higher levels compared to *WT* (Fig. 3I, S12A-B), consistent with our findings. Given the disappearance of colon CXCR4^+^ B cells and the presence of such cells within the CNS in the *5XFAD* mice, we next determined if any CNS infiltrating B cells and/or ASCs are potentially linked to the gut. Thus, we performed a gut commensal ELISPOT assay, which identifies ASCs that recognize gut-derived commensal antigens [27] in the dura mater of mice. Remarkably, we found a significant increase in IgA^+^ and a near significant increase in IgG^+^ gut-specific ASCs in the *5XFAD* dura mater (Fig. 3J). Glial cell induction of CXCL12 and brain enrichment of CXCR4^+^ B cells and IgA^+^ cells in the dura may implicate the action of this chemokine on the gut-brain axis.

### Inulin fiber improves features of AD and associated gut and brain parameters in *5XFAD* mice

Given the increased accumulation of DN B cells and IgA^+^ ASCs expressing gut migratory markers in the brain, coupled to a reduction of CXCR4^+^ B cells in the colon of *5XFAD* mice, we next tried to target this axis by boosting potentially protectively gut IgA^+^ ASC levels in the colon. During homeostasis, intestinal IgA^+^ ASCs produce IL-10 in addition to opsonizing commensal antigens, acting to maintain microbial homeostasis in the gut [78]. While IgA ASC migration into the CNS is thought to attenuate neuroinflammation [27,28], its loss in the colon might promote gut dysbiosis and an eventual loss of regulatory IgA responses. We, therefore, predicted that boosting IgA responses in the gut would simultaneously reverse the leaky gut phenotype predicted in AD intestines [32] and dampen neuroinflammation in AD. Fiber diet interventions, particularly inulin, boost systemic T_regs_ levels via the gut microbiome-derived SCFAs [79]. In turn, T_regs_ are thought to boost IgA levels via their action on B cell isotype switching [80,81]. Inulin also promotes type 2 immune responses via bile acid production [35].

First, we investigated tolerogenic transcriptional signatures in T_regs_ in the *5XFAD* colon using our scRNA sequencing data. The thymus-derived HELIOS^+^FOXP3^+^ T_reg_ subcluster showed lower levels of *Il10*, *Tgfb*, and *Ctla4* mRNA, whereas microbiome-derived HELIOS^-^ T_regs_ were transcriptionally unchanged compared to *WT* [82] (Fig. S13A). While we found no major changes in total T cells or T_regs_ within the colon by flow cytometry (Fig. S13B-C), T_reg_ frequencies were markedly reduced in the spleen, brain, and blood (Fig. S13D-F). Interestingly, colon T_regs_ were partially CXCR4^+^, and these cells showed a significant reduction in *5XFAD* mice (Fig. S13G), much like earlier observed in ASCs. These T_regs_ may also migrate towards a CXCL12 gradient in the brain and attenuate neuroinflammatory gliosis [83,84]. Assessment of microbiome-derived T_regs_ is discussed below.

To test the effect of fiber diet interventions on AD pathology, we fed mice with control (4.7% cellulose), cellulose (17%), or inulin (17%)-enriched diets from 8 weeks of age till the end of the study at 15 months (Schematic: Fig. 4A). An enlarged cecum in the inulin-fed mice, compared to both cellulose and control diets, was consistent with microbial metabolism of these fibers (Fig. S14A). Compared to control and cellulose-fed mice, inulin-fed mice showed significant improvements in frailty in both *WT* and *5XFAD* groups, including marked changes in body weight (Fig. 4B-C, S14B). Interestingly, cellulose fiber led to a trending worsened age-related frailty in *WT* mice but protected against frailty in the *5XFAD* group (Fig. 4C, S14B). These changes were mirrored in the open-field behavioral assay with undetectable changes in the Y-maze assay (Fig. S14C). Quantification of Aβ plaques in the hippocampus and cortex showed that inulin and cellulose diets resulted in a reduction of Aβ plaques and correlated with reduced immune cell (CD45^+^) infiltration (Fig. 4D). In inulin-, but not cellulose-fed mice, we observed an induction of IgA^+^ cells in both *WT* and *5XFAD* conditions in both the colon and the spleen (Fig. 4E-F). These IgA^+^ cells were CXCR4^+^ (Fig. S14D). We also observed an induction of gut T_regs_ likely driven by increased SCFAs expected to be generated by an inulin-enriched microbiome (Fig. 4G) [85]. The numbers of RORγt^+^ T_regs_, representing the gut-specific pT_reg_ population were reduced in control diet *5XFAD* colons (Fig. 4H). The numbers of colon pT_regs_ were increased in inulin and not cellulose-fed mice, rescuing the loss in *5XFAD* (Fig. 4H). The significant reduction of T_reg_ frequencies in the *5XFAD* spleen was also rescued under inulin diet conditions (Fig. 4F). Interestingly, the inulin diet also resulted in reduced IBA1 and CXCL12 levels in the brain (Fig. 4I). Thus, the inulin diet may be one intervention that can boost regulatory immunity within the intestine while reducing accumulation of microglia and their chemokine axes within the brain, alleviating Aβ deposition and some neurodegenerative manifestations of AD.

**Figure 4:**
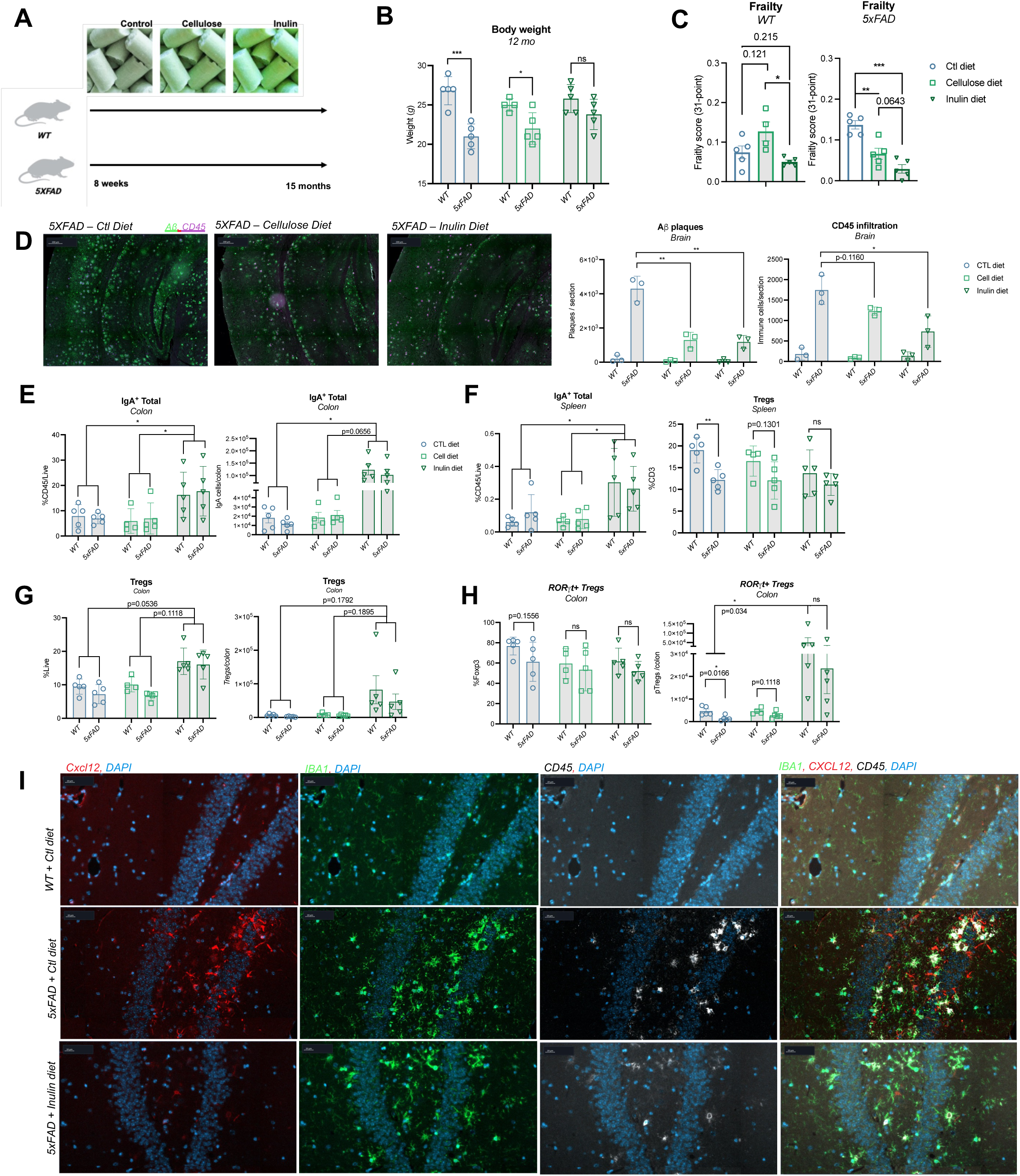
Inulin fiber rescues AD parameters while boosting colon IgA and T_reg_ levels and dampening gliosis associated CXCL12 in the brain. **(A)** Schematic of long-term dietary intervention study. **(B)** Body weight of *WT* and *5xFAD* 12-month-old female mice at on control (circle), cellulose (square), and inulin diets (triangle); n= 4-5 per group. **(C)** 31-point frailty index score of mice at 12 months. **(D)** Representative immunofluorescence staining of FFPE hippocampi from *5xFAD* mice on each specified diet using CD45 (purple) and Aβ (6E10) (green) antibodies (left) at endpoint. Quantification of extracellular Aβ plaques and cellular CD45 staining in each section; n=3 per group (right). **(E-G)** Flow cytometric quantification of colonic immune cell frequencies of IgA^+^, total Foxp3 ^+^ T_regs_ and total Foxp3^+^Rorgt^+^ pT_regs_ at endpoint; n=4-5 per group. **(H)** Flow cytometric quantification of splenic immune cell frequencies of total IgA^+^ and Foxp3^+^ T_regs_ at endpoint; n=4-5 per group. **(I)** Representative immunofluorescence staining of FFPE hippocampi of *WT* control diet (top) and *5xFAD* control diet (middle) and *5xFAD* inulin diet (bottom) using IBA1 (green), CXCL12 (red), and CD45 (white) antibodies.

## Discussion

Changes in gut immunity and microbial homeostasis have been reported in neuroinflammation and brain injury [21,22,86]. However, the nature of gut-brain crosstalk in AD is not well understood. Here, we examined the gut immune landscape in AD using a pre-clinical Aβ-AD model, the *5XFAD* mouse. At the global level, we found pan-immune transcriptional changes in the colon, including in genes involved in ribosomal activity, mitochondrial stress, and immune activation. These gut immune changes were not associated with any accumulation of Aβ in the *5XFAD* colon. Specifically in B lineage cells of the colon, we saw a striking loss of CXCR4^hi^ ASCs and increased B2 class switching, collectively reflecting gut B cell dysfunction in the context of AD. Because humoral immunity is a key player in maintaining gut microbial homeostasis [87], we suspect these B lineage changes are closely linked to reported microbiome shifts [88,89] and leaky gut phenotypes in Aβ-AD mice [31,32] and suspected in AD patients [18–20].

While the migration of gut ASCs into the brain and brain border regions is emerging as a novel component modulating neurological disease [27,28], its potential involvement in AD has not been fully explored. In our colony of *5XFAD* mice, we found a striking loss of colonic ASCs with a migratory signature. This migratory signature suggests an ability to respond to various signals, including CXCR4/7 ligands, S1P, BAFFR ligands, FFAR1 binding medium-chain fatty acids, and GPR183-binding oxysterols. Some of these have been linked to resilience in AD. For example, GPR183-mediated migration [90] may be affected by APOE2 oxysterol production in the brain [91]. Here, we focused on CXCR4 as we observed downregulation of CXCR7 (*Ackr3*) in gut ASCs, a factor that increases CXCR4 responsiveness to CXCL12 in B cells [89]. Furthermore, strong CXCR4 expression has been reported on IgA^+^ cells found in the meninges [28].

Within the *5XFAD* brain, we found a significant increase in CXCL12 levels, the cognate binding partner for CXCR4. Increased levels of CXCL12 were produced by AD microglia and astrocytes, which accumulated at the hippocampal formation, representing neuroinflammatory gliosis. The accompanying reduction in endothelial CXCL12 may reflect the reported degeneration of the blood-CSF barrier in AD [92] and mirror CXCL12 polarization-mediated trafficking of cells into the brain parenchyma as suspected in MS [93]. Already recognized as a key mediator of AD-associated neuroinflammation, gliosis may also activate a CXCL12-CXCR4 migratory axis signaling gut ASC migration into the brain and border regions. Using commensal ELISPOT assays, we confirmed increased levels of gut-specific IgA^+^ ASCs in the AD dura mater. We also found increased CXCR4^+^ B cells within the AD brain parenchyma. CNS-infiltrating B cells have been linked to AD pathophysiology [94]. The role of gut IgA ASCs in the CNS is likely multifactorial, including potential protective or compensatory effects on inflammation [27,28]. The CXCL12-CXCR4 axis has been implicated in B cell development [73] and ASC trafficking towards inflammatory sites [95,96], as well as in facilitating crosstalk between dura fibroblasts and immature B cell progenitors in the skull bone marrow [74,75]. Additionally, we found increased CXCR4 expression in other immune cell subtypes, suggesting that CXCL12-mediated recruitment is not likely limited to gut-derived ASCs. While CXCL12 expression has been observed in the colon [97] and bone marrow [96], we expect heightened glial cell expression of CXCL12 to partially redirect CXCR4-trafficking, including of circulating gut immune cells, into the AD brain parenchyma. Interestingly, changes in CXCL12 expression in α-synuclein-mediated Parkinson’s disease [98] and MS [93] have been reported, suggesting this axis may be a key feature shared by several neuroinflammatory conditions.

Due to the identification of gut commensal-responsive cells in broad CNS tissues, our study warrants further experimentation using small-molecule CXCR4 inhibitors (e.g., AMD3100 [99]), CXCR7 blockade [100], or CXCL12 targeting (e.g., NOX-A12 [101,102]) to better dissect gut-brain migration. Such studies may offer a method to block gut-brain ASC migration and determine its neuroprotective role while informing on potential therapeutic targets and adverse reactions.

Another key finding of our work is the presence of multiple markers of potentially low-grade systemic inflammation within the blood and spleen of *5XFAD* mice. We found increased B2 cell class switching throughout the periphery and a systemic reduction of T_reg_ levels at the investigated times (9-12 months of age). A similar B cell phenotype has been reported in human PBMC studies, implicating a change in BCR repertoire with AD pathology [70]. While some BCRs are expected to generate Aβ auto-antibodies [103], it remains to be seen whether these BCRs are specific to gut microbial antigens, including microbial amyloids [104] in AD, accumulating in the periphery due to a leaky gut [20,31,32].

To determine whether rescuing inflammatory phenotypes within the AD gut could constitute a potential therapeutic, we looked at intervention utilizing the dietary supplement, inulin. Inulin is a soluble prebiotic fiber that alters systemic levels of several metabolites, including SCFAs [105], and decreases neuroinflammation in other mouse models of AD [106]. In our model, consumption of dietary inulin resulted in protection against various hallmarks of AD. Upon inulin feeding, we found a systemic increase in T_regs_ levels in *5XFAD* mice and restored pT_reg_ levels in the colon, predicted to be driven by SCFA production [79]. Additionally, we found increased gut IgA^+^ cell levels in inulin-fed mice, likely driven by T_reg_-mediated effects on isotype switching [81,107,108]. These results suggest that microbial fermentation of inulin leads to modulation of gut T_regs_ and IgA^+^ cells. These IgA^+^ cells are CXCR4^+^ that, as alluded to earlier, may migrate towards CXCL12 signals from AD brain gliosis. Interestingly, we also observed reduced glial CXCL12 production upon inulin feeding, predicted to be at least partially driven by systemic SCFA accumulation [23,24]. Shown to also act on microglia in the brain, SCFAs accumulating in the periphery may act directly on brain microglia via SCFA transporters [24,109] and TREM2 activation [110]. The impact of SCFAs or other inulin-derived metabolites on astrocyte CXCL12 expression is unknown. The anti-inflammatory effects of inulin-derived metabolites may also include gut epithelial remodeling [111], induction of regulatory ILC2s in the gut [35], and protection via brain T_regs_. The protection conferred by inulin in the gut is also predicted to attenuate systemic inflammation driven by leaky gut antigens [111]. Overall, the inulin diet likely rescued against AD hallmarks via multiple mechanisms acting on the gut, the brain, and peripheral immune cells.

Our data support a model whereby AD facilitates gut and systemic inflammatory alterations in association with increased CXCL12 production from CNS resident microglia and astrocytes. More work is needed to determine the relative contributions of local versus systemic and gut-related inducers of glial CXCL12. Increased CXCL12 in the CNS, in turn, may promote migration of immune cells, including B cells with inflammatory capacity, as well as IgA^+^ gut ASCs with the potential to dampen or promote inflammation. The reduction of B cell subsets from the colon potentially compromises local intestinal defenses, facilitating reported gut microbial changes linked to AD progression [32,69,88]. These processes are targetable by dietary supplements, including soluble fiber that inhibits the chemokine cascade and restores gut and systemic immune regulatory cells resulting in the dampening of AD disease progression. Our investigation provides a better understanding of colon immune responses in AD and reveals key links to neuroinflammation. Through this investigation, we provide further evidence for the importance of the gut-brain axis, including the role of gut immune cells and reveal possible dietary, and therapeutic interventions to target inflammation in AD.

### Limitations of the study

Although gut ASCs are expected to attenuate inflammatory responses to commensal species within the gut, the protective nature of brain and dura mater migrant gut ASCs remains unclear. In autoimmunity, IgA^+^ ASCs produce IL-10 to attenuate neuroinflammation [27]. In AD, IVIg delivered with a BBB-penetrating treatment suggests that antibodies have a protective effect [112]. In contrast, depletion of immature and mature B cells using biologics appears to be protective in neurological conditions, including AD [94,113]. Temporal activation signals acting differentially on heterogeneous B cell subsets may account for these observations. The BCR repertoire in gut ASCs may also drive differential inflammatory effects upon brain migration. For example, gut IgA^+^ ASCs have isoform-dependent heterogeneity in the engagement of Fcα receptors that can differentially modulate inflammation in humans [114]. This implicates, for example, *Trichomonas* levels [115] and previous *Salmonella* encounters [116] in certain individuals and their baseline IgA repertoire. The role of BCRs against microbial amyloids [104], as well as brain Aβ, encountered via gut-CSF encounters [106] is also poorly understood. Studying brain migrating gut ASC repertoires would allow determination of whether and which ASCs are protective.

The route of migration of gut immune cells into the brain in AD remains enigmatic. Possible routes include the BBB, the meningeal lymphatics, or the vagus nerve [117]. Although we find trending increases in DN B cells and IgA^+^ cells in *5XFAD* blood, it is unclear whether these are mainly driven by the systemic accumulation of gut bacterial antigens via a leaky gut [31,32], or by their migration via the bloodstream [118] or some combination of the two. Gut-specific IgA^+^ ASCs within the dura mater imply that lymph-CSF barrier dysfunction might be at play, already associated with AD [119]. Immune cells within human AD CSF also show reduced CXCR4 expression [120], suggesting possible infiltration into tissue, though early reductions in CSF CXCL12 levels in AD suggest a temporal effect [121].

Here, we only studied the Aβ component of the disease with single-cell analysis from a single mouse model of AD. Given the importance of the gut microbiota in AD, it will be important to validate gut immunological signatures across different models and in different facilities. Consistently, because immune responses to the extracellular Aβ and the intracellular tau are expected to be different [122], it is unclear whether these phenotypes are relevant to tau-driven late stages of AD. Finally, though we have provided evidence for some of our conclusions through use of publicly available human data, a similar immunophenotyping of matched human brain and colon tissue would be required to improve the translational conclusions of our work.

## Acknowledgments

We thank the Buck Institute flow cytometry core facility for their assistance with high-dimensional spectral flow cytometry, specifically Dr. Herbert Kesler, Dr. Ritesh Tiwari and Ryan Kwok. Graphical illustrations were made via BioRender.com. This work was mainly supported through funds derived from the National Institutes of Health (NIH) grant 3RF1 AG062280-01S1 (J.K.A and D.A.W.). This work was also supported in part via the Canadian Institutes of Health Research (CIHR) grant PJT186165 (D.A.W.) and in part through funds derived from the Buck Institute for Research on Aging (D.A.W). K.A.W and T.R.V. were supported via the T32 NIH fellowship grant NIA T32 AG000266 with L.M.E and D.A.W., respectively. Funding support for L.M.E. was from NIH grant, AG066591. O.L.R. was supported via the startup grant, Krembil Research Institute 410013711.

## Author Contributions

P.M., O.L.R., J.K.A, L.M.E. and D.A.W. contributed to the experimental design, execution, analysis, and writing of the manuscript for this study. P.M., R.E., A.R., J.H.Y.N., T.R.V., C.R.T, H.D., and K.A.W. conducted *in vivo* experimentation. P.M., C.G.A, W.C.M., and F.W. performed computational analyses of single cell datasets. M.M.M. performed analysis of immunofluorescence microscopy images. S.K., L.M.E., K.A.W., C.W., A.M. and S.W. provided experimental feedback and/or manuscript editing support. D.A.W, J.K.A., O.L.R., L.M.E., and D.F. supervised aspects of the study. D.A.W., O.L.R. and J.K.A. supervised the overall implementation of the study. All authors had an opportunity to view and edit the manuscript.

## Competing Interests

D.A.W. is a co-founder of Propion Inc., a company that studies gut immune interventions for aging and related diseases. Other authors of this manuscript declare no competing interests.

