## Supplementary Figure Legends for "Single-cell transcriptomics reveals colonic immune perturbations during amyloid-β driven Alzheimer’s disease in mice"

**Figure. S1 (related to Figure 1): Evaluating systemic changes in *5xFAD* versus *WT* mice. (A)** 31-point frailty index score per category *5xFAD* and *WT* female mice at 12 months of age, n = 5 per group. **(B)** Colon length, spleen weight and colon length to weight ratio in *5xFAD* (square) to *WT* (circle) female mice at 12 months of age; n=5-10 per group. **(C)** Open field behavioral assay measurements in female mice at 12 months. **(D)** Small y-maze behavioral assay measurements in female mice at 12 months. **(E)** H&E staining of colon Swiss rolls from 6-month and 12-month-old female mice. **(F)** IF staining of spleens from 9-month-old female mice with B220 (red) and CD4 (yellow) (left) and normalized follicle measurements (right).

**Figure S2: (related to Fig. 1) Evaluating systemic Aβ accumulation.** Colon (top) and brain (bottom) sections stained with CD19 (red) and Aβ (clone 6E10) (green) antibodies from 12-month-old *WT* (left) and *5xFAD* (right) female mice.

**Figure S3 (related to Fig. 1): Capturing cluster specific heterogeneity and differential gene expression in colon immune cells. (A)** Heatmap of top 50 genes per cluster. **(B)** Heatmaps of a curated set of genes representing known genes that capture B cell (top) and T/ILC (bottom) heterogeneity. **(C)** Top DEGs per cluster; (adj. p) cutoff is 0.05 and log2FC cutoff is 0.25.

**Figure. S4 (related to Fig. 1): Trajectory analysis of B cell subclusters using *Monocle3*. (A)** UMAP of all possible trajectories of *WT* and *5xFAD* colon B cells. Black circles indicate branch nodes, where cells can travel to a variety of outcomes. Light gray circles designate different trajectory outcomes. **(B)** Gene expression changes by specified B cell development gene, across pseudo time starting at naive B cells (cluster 0).

**Figure S5 (related to Fig. 2): Representative gating strategy for the flow cytometric assessment of colon immune cells.**

**Figure S6: (related to Fig. 2): Flow cytometric assessment of colon immune cells (A)** Representative flow cytometry plots of intracellular BLIMP1 expression and gating in colonic CD45^+^CD3^-^ cells in each genotype (left). Frequencies and cell numbers of total BLIMP1^+^ cells in the colon (right). Female mice at 12 months old from *WT* (circle) and *5xFAD* (square) were used; n=5 per group. **(B)** Frequencies and cell numbers of total CD45^+^IgA^+^ cells in the colon measured by flow cytometry. **(C)** Immunofluorescence staining of FFPE colon swiss rolls with intracellular IgA (orange) antibody in the lamina propria of 12-month-old female *WT* (left) and *5xFAD* mice (right). **(D)** Representative flow cytometric plots of CXCR4 expression within the total B220CD43 DN B cell subset (left) and cell numbers of CXCR4^+^ cells within each DN isotype subgroup (right). **(E)** Representative flow cytometric plots of JUN expression within the B220CD43 DN B cell subset (left) and cell numbers of JUN^+^ cells within each DN isotype (right). **(F)** Immunofluorescence staining of FFPE colon swiss rolls with JUN (orange) and CD19 (green) antibodies highlighting JUN^+^ B cells (arrows) in proximity to isolated lymphoid follicles in 12 month female *WT* (left) and *5xFAD* mice (right). **(G)** Frequency and cell number of CD19^+^ B cell subsets including B220SP, CD43SP and B220^-^CD43^-^ DN cells (left) and cell numbers of each isotype (IgA, IgM, or triple negative with IgD) within the B220CD43 DN B cell population (right). **(H)** Frequency and cell number of each immune cell subtype within the colon including CD19^+^ B cells, CD3^+^ T cells, CD127^+^ ILC, and CD11b and CD11c expressing myeloid cells.

**Figure S7 (related to Fig. 2): Transcription of selected genes in colon B cells including ribosomal, mitochondrial and immune process genes.** Violin plots generated using SCTransform normalized scRNAseq data from *WT* and *5xFAD* mouse colon B cells.

**Figure S8 (related to Fig. 2): Representative gating strategy for the flow cytometric assessment of peripheral immune cells. (A)** Spleen **(B)** Blood**.**

**Figure S9 (related to Fig. 2): Flow cytometric analysis of peripheral immune cells (A)** Representative flow cytometric plots of splenic CD19^+^ B cell subsets (left) and quantified frequencies (right) in 12 month old *WT* (circle) and *5xFAD* (square) female mice; n= 4-5 per group. **(B)** Representative plots of blood CD19^+^ B cell subsets (left) and quantified frequencies (right) in *WT* (circle) and *5xFAD* (square) female mice; n= 5 per group. **(C)** Frequencies of blood (left) and spleen (right) total IgA^+^ cells in 12mo *WT* (circle) and *5xFAD* (square) female mice; n= 5 per group. **(D)** Representative plots of splenic B220SP^+^ B2 cell CD86 and MHCII expression (left) and quantified frequencies of CD86^+^ B2 cells (right) in *WT* (circle) and *5xFAD* (square) female mice; n= 4-5 per group. **(E)** MHCII expression (gMFI) in blood B cells in 12mo *WT* (circle) and *5xFAD* (square) female mice; n= 5 per group. **(F)** Representative plots of flow cytometric assessment of splenic B220SP^+^ cell subsets by CD23 and CD21 denoting marginal zone B cells (MZB); follicular B cell (FOB) or age-associated B cells (ABCs) (left). Frequencies of each subset (right) in *WT* (circle) and *5xFAD* (square) female mice; n= 5 per group.

**Figure S10 (related to Fig. 3): Representative gating strategy for the flow cytometric assessment of brain and brain associated immune cells. (A)** Brain parenchyma **(B)** Dura mater.

**Figure S11 (related to Fig. 3): Flow cytometric analysis of brain immune cell infiltration. (A)** Representative plots showing CD45^med^, CD45^hi^ and CD45 total immune cells within the brain parenchyma (left) quantified by frequency and cell number in *WT* (circle) and *5xFAD* (square) female mice at 12mo (right); n= 5 per group. **(B)** Representative gating strategy (left) and frequencies of total CD19^+^ B and CD3^+^ T cells in the CD45^hi^ lymphocyte population (right); n= 5 per group. **(C)** Representative plots showing the CD45^-^CD31^+^ endothelial cell population in the brain parenchyma (left) and quantified within the *WT* (circle) and *5xFAD* (square) groups. **(D)** Representative histogram gating strategy for CXCL12 expression in brain endothelial cells (left) and frequencies (right) within the *WT* (circle) and *5xFAD* (square) groups.

**Figure S12 (related to Fig. 3): Mining publicly available brain CD45 immune cell scRNAseq data from *5xFAD* mice***.* **(A)** UMAP of reanalyzed mouse brain CD45^+^ scRNAseq from Keren-shaul *et* *al.* 2017 in *WT* and *5xFAD* mice (top) showing *Cxcr4* transcript appearing in granulocyte and T/B cell clusters (bottom). **(B)** UMAP of reanalyzed mouse brain CD45^+^ scRNAseq analysis from Su *et* *al.* 2023 in *WT* and *5xFAD* mice with CD19 B and CD3 T clusters highlighted (top). Migratory gene transcripts of *Cxcr4*, *Klf2*, *Jun* and *S1pr1* in *WT* and *5xFAD* mice (bottom).

**Figure S13 (related to Fig. 4): Regulatory Treg status in mouse colons. (A)** UMAP of T/ILC clusters derived from colon immune cells in *WT* and *5xFAD* mice showing *Foxp3*, *Il10*, *Ctla4*, *Helios*, *Tgfb1*, and *Cxcr4* expression patterns. **(B)** Representative flow cytometric gating strategy for total colon T cells (left) with quantification of frequency and cell number per genotype at 12 months; n=5 per group. **(C)** Representative flow cytometric gating strategy for total colon FOXP3^+^ T_regs_ (left) with frequency and cell number per genotype (right); n =5 per group. **(D)** Representative flow cytometric gating strategy for splenic T cells and T_regs_ (left) and quantification of T_reg_ frequency and cell number in 12-month *WT* and *5xFAD* female mice (right); n =5 per group. This was repeated for **(E)** brain and for **(F)** blood. **(G)** CXCR4 gMFI within the FOXP3^+^ T_reg_ population in the colon by genotype.

**Figure S14 (related to Fig. 4): Measuring AD parameters in fiber diet fed mice (A)** Representative cecum images of control diet (left) and inulin diet (right) mice at endpoint. **(B)** 31-point frailty index score per category in *WT* (top) and *5xFAD* (bottom) female mice at 15 months of age after dietary intervention with control (circle), cellulose (square) or inulin (triangle) fibers, n = 4-5 per group. **(C)** Open field (left) and small y-maze (right) behavioural assay measurements in female mice at 15 months. **(D)** Flow cytometric assessment of colonic CXCR4^+^IgA^+^ ASCs quantified by frequency (left) and cell number (right) in 15-month-old diet fed mice as indicated.
