## Supplementary figures and images for "Single-cell transcriptomics reveals colonic immune perturbations during amyloid-β driven Alzheimer’s disease in mice"

### Fig. S1

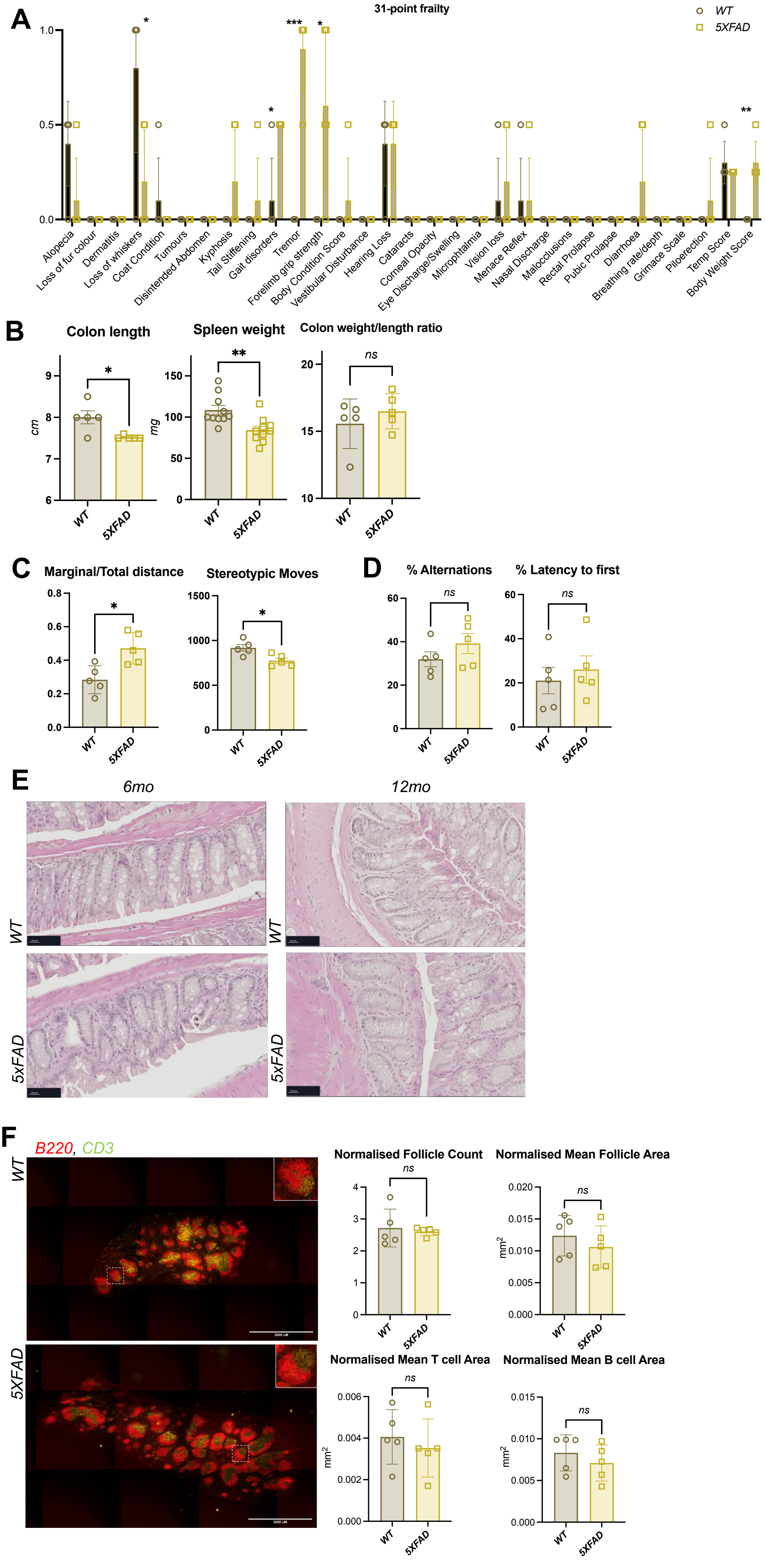

### Fig. S2

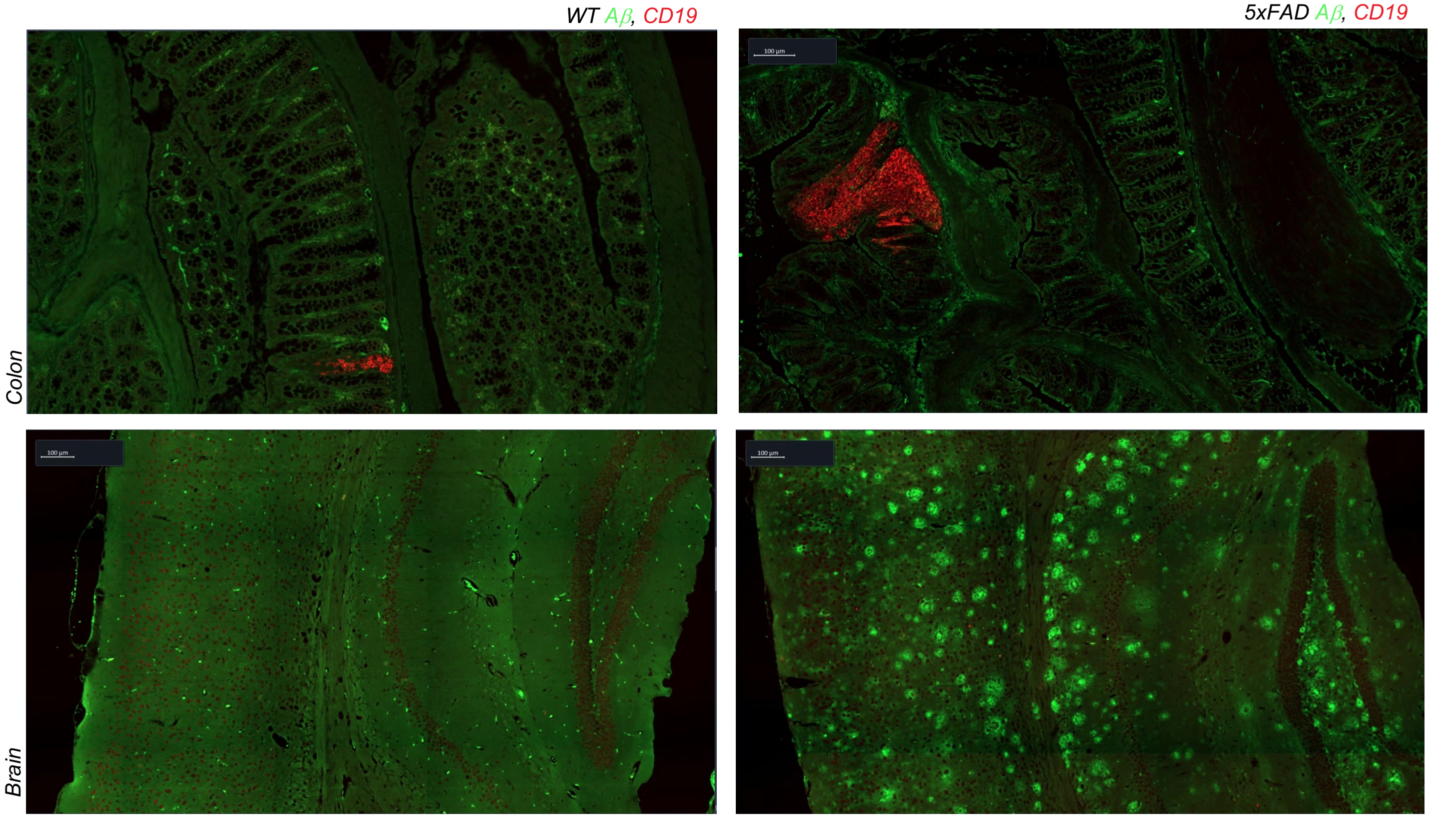

### Fig. S3

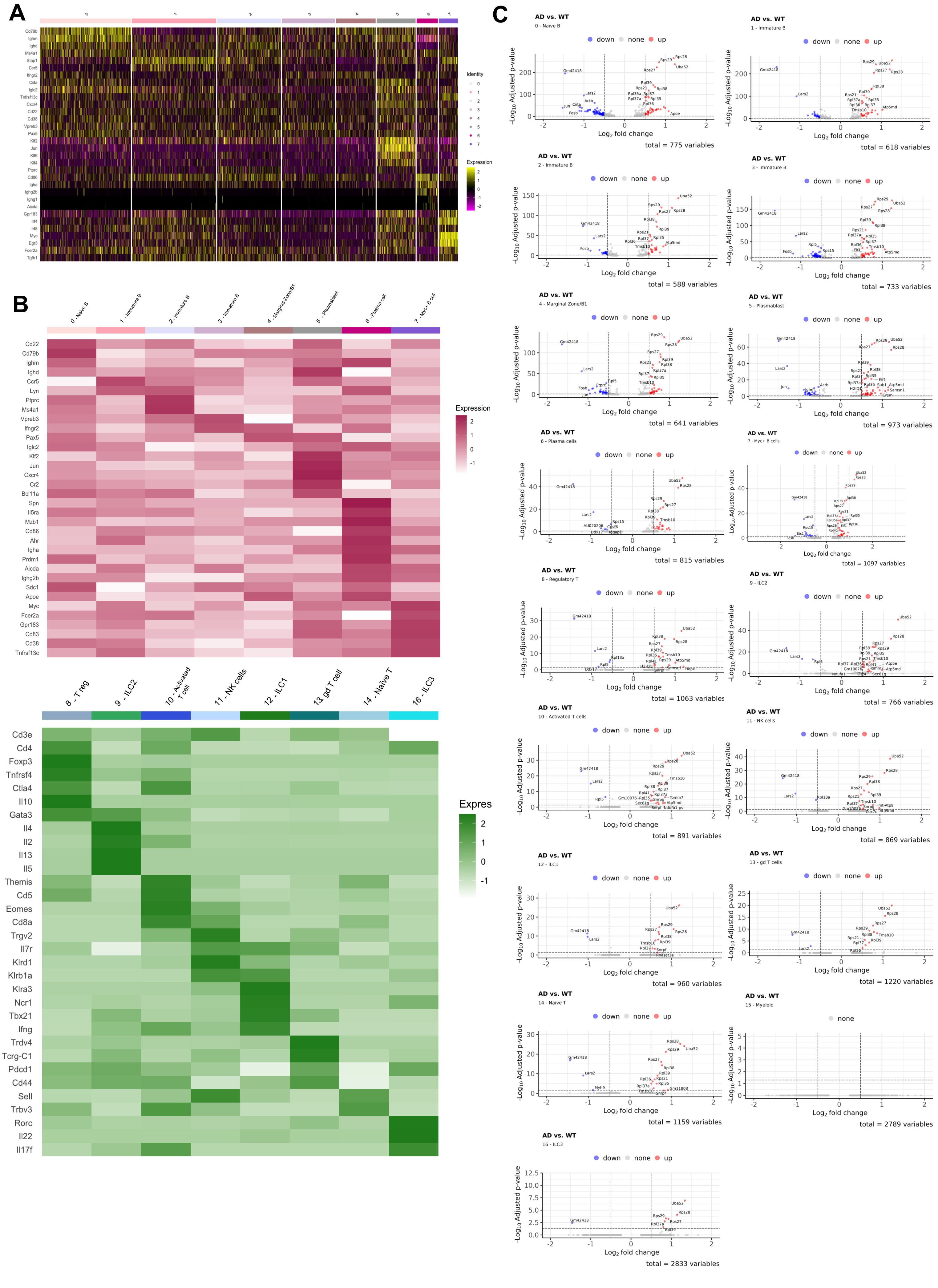

### Fig. S4

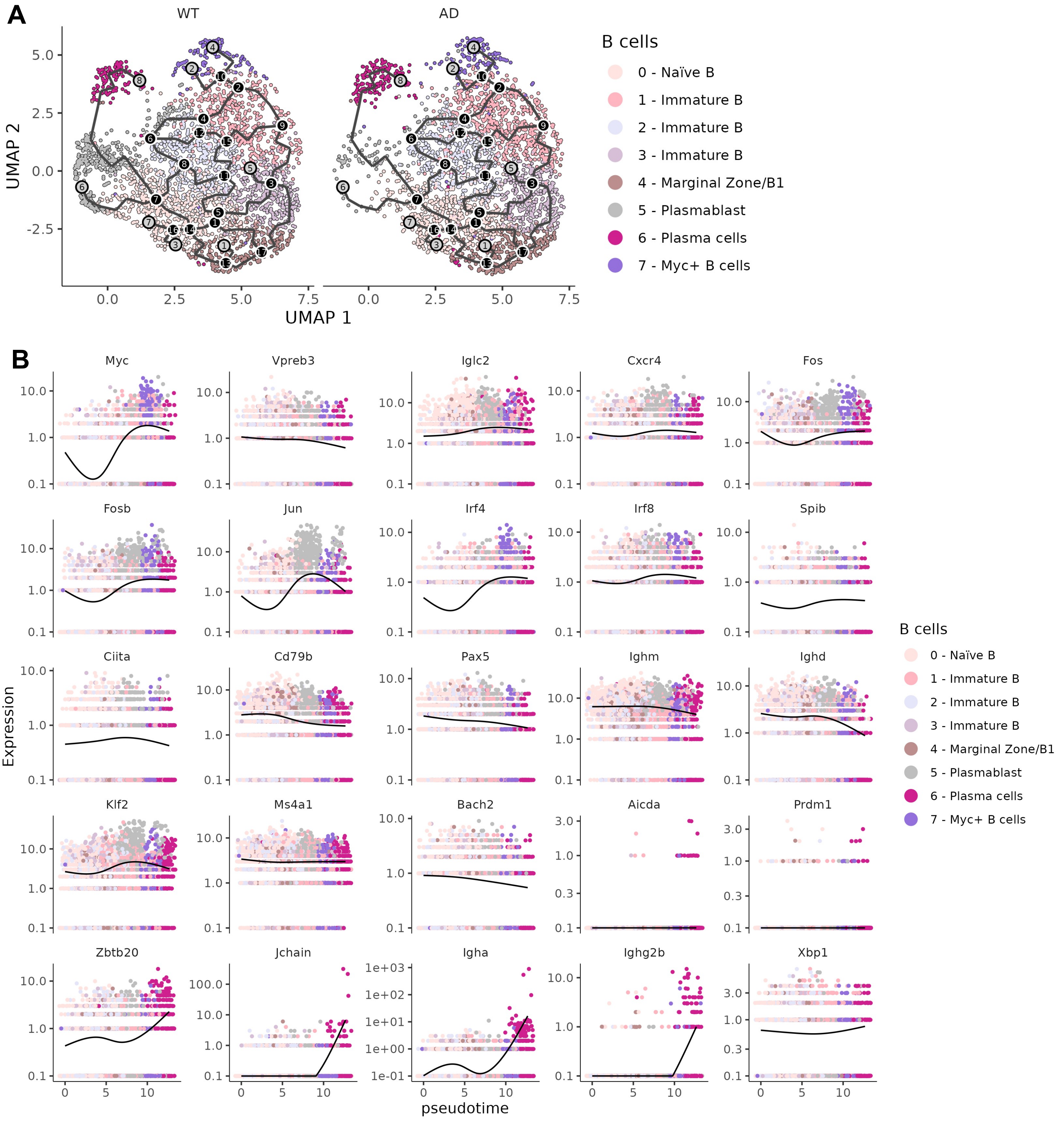

### Fig. S5

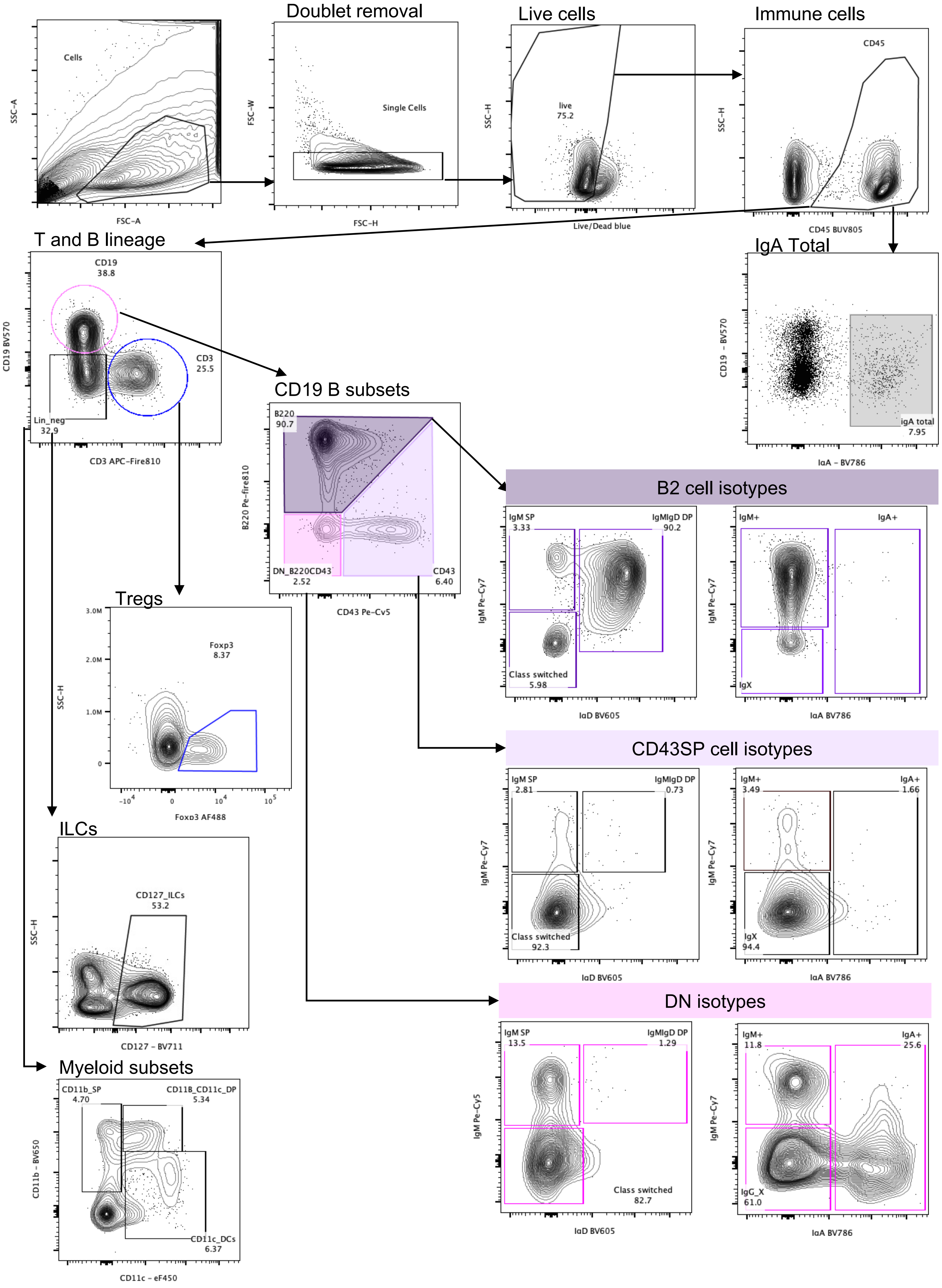

### Fig. S6

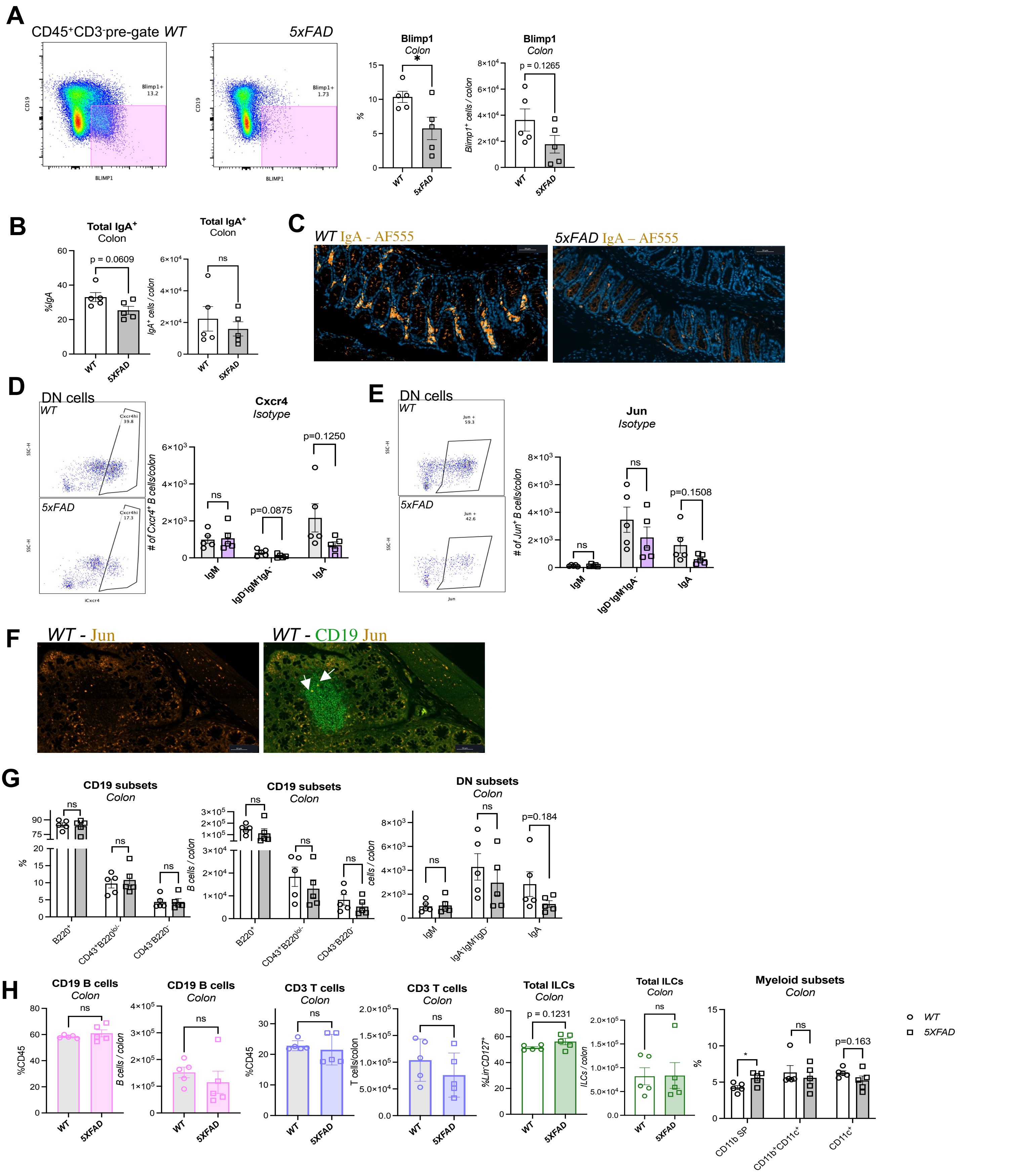

### Fig. S7

#### Selected key biological process genes across B cell lineages

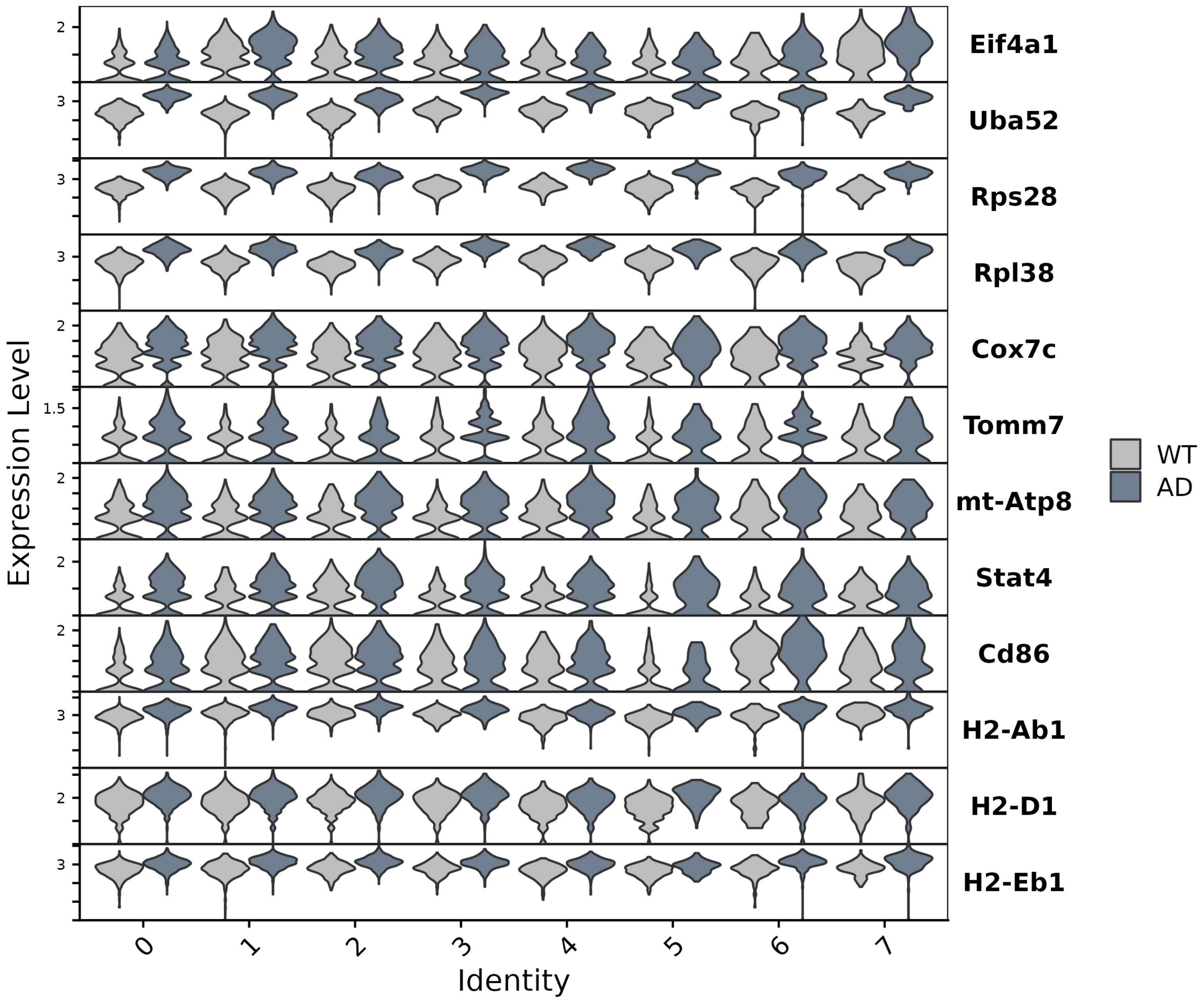

### Fig. S8

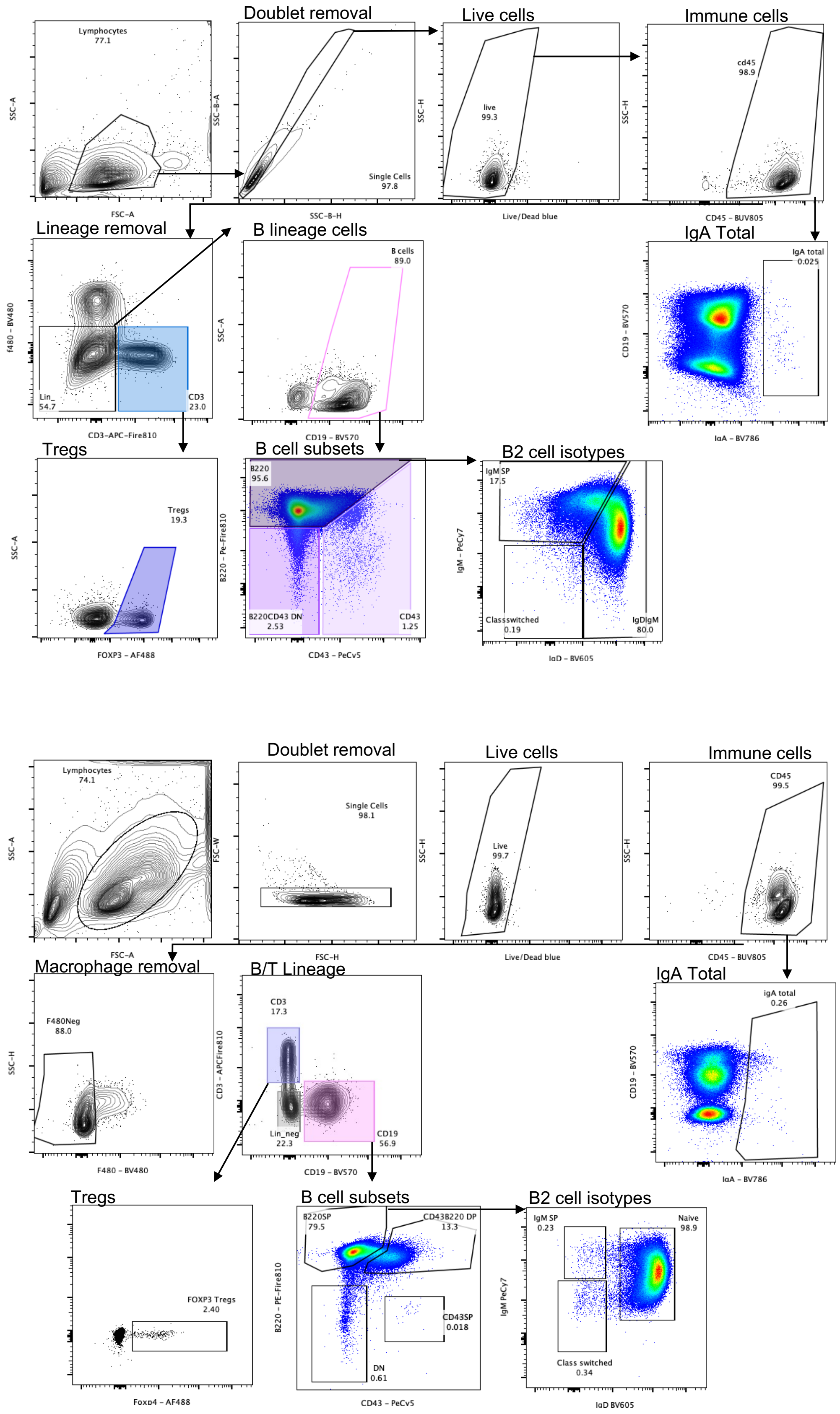

### Fig. S9

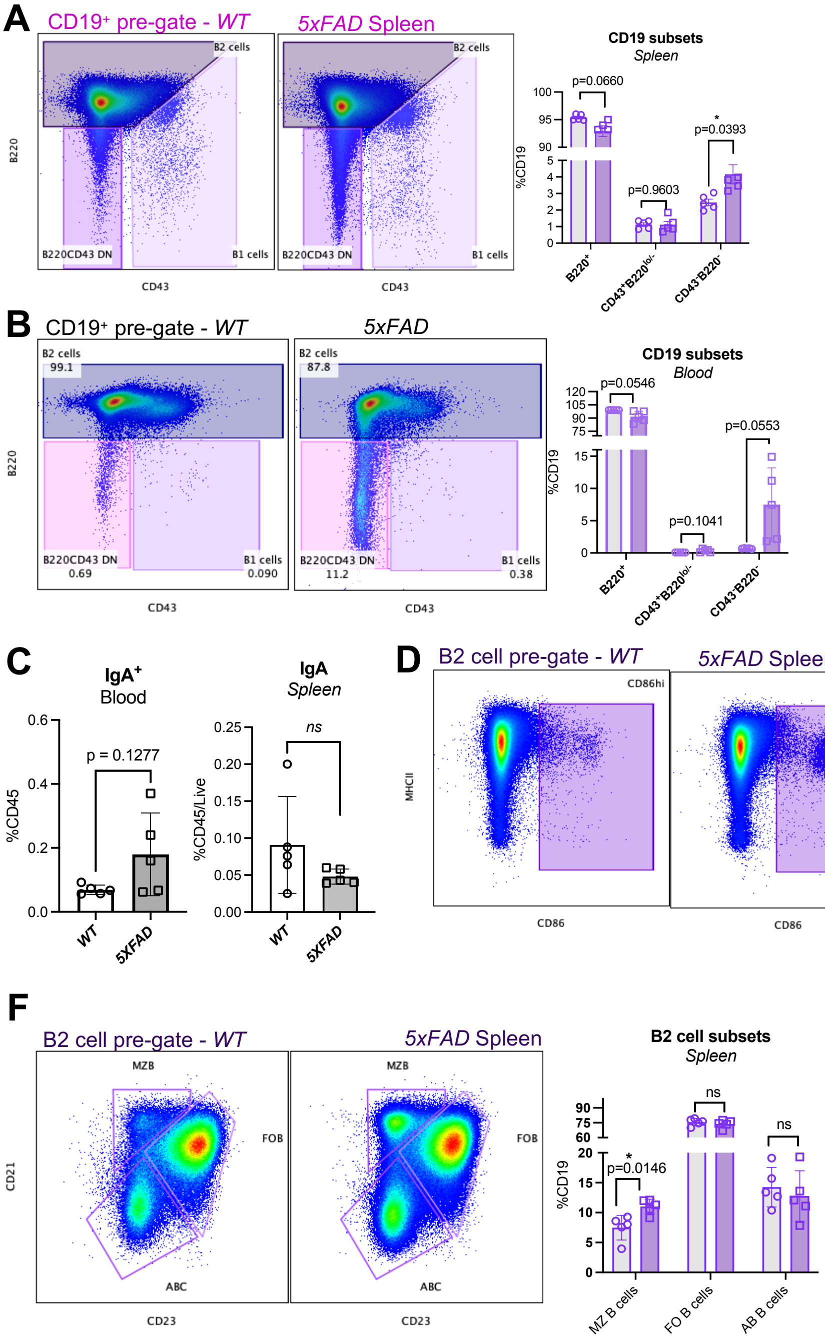

### Fig. S10

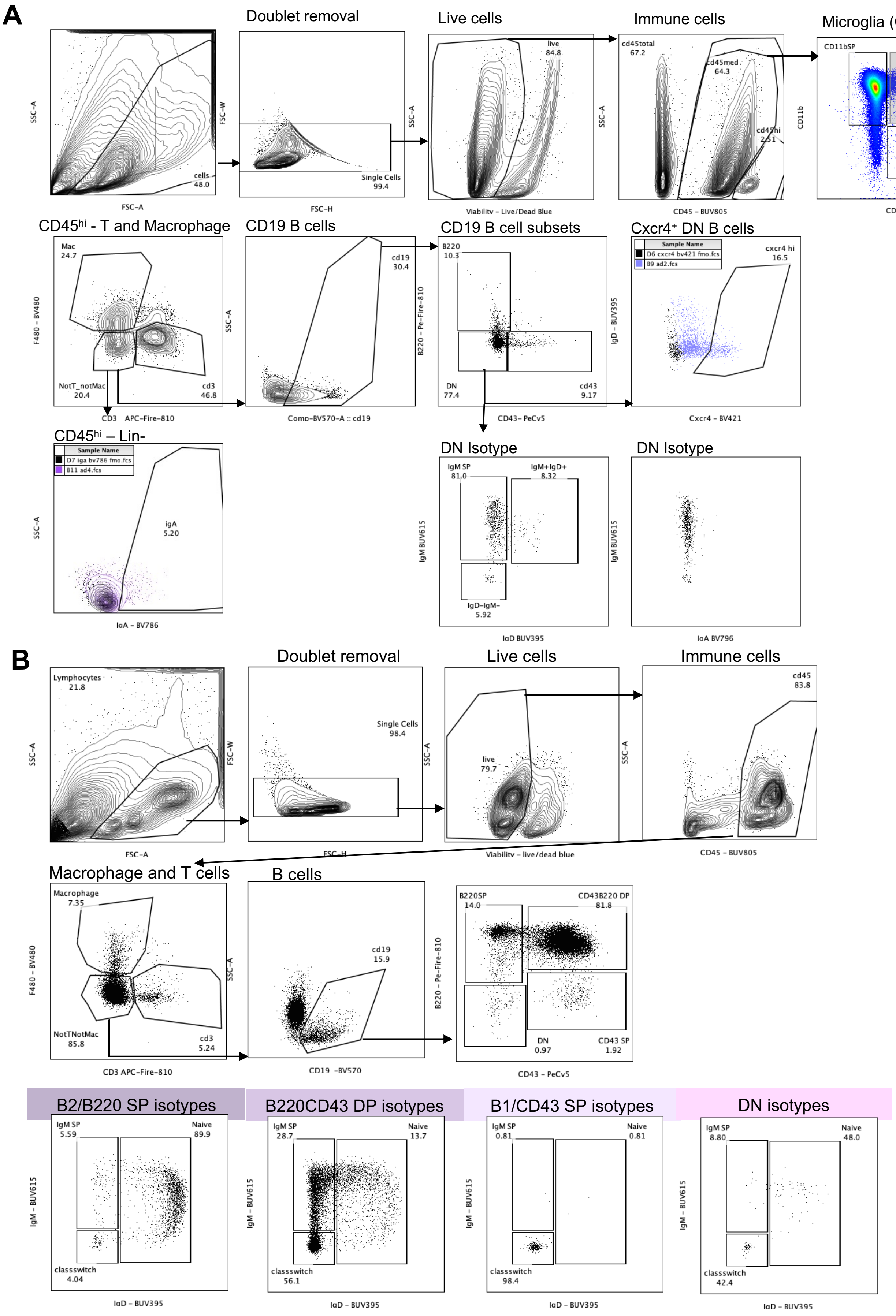

### Fig. S11

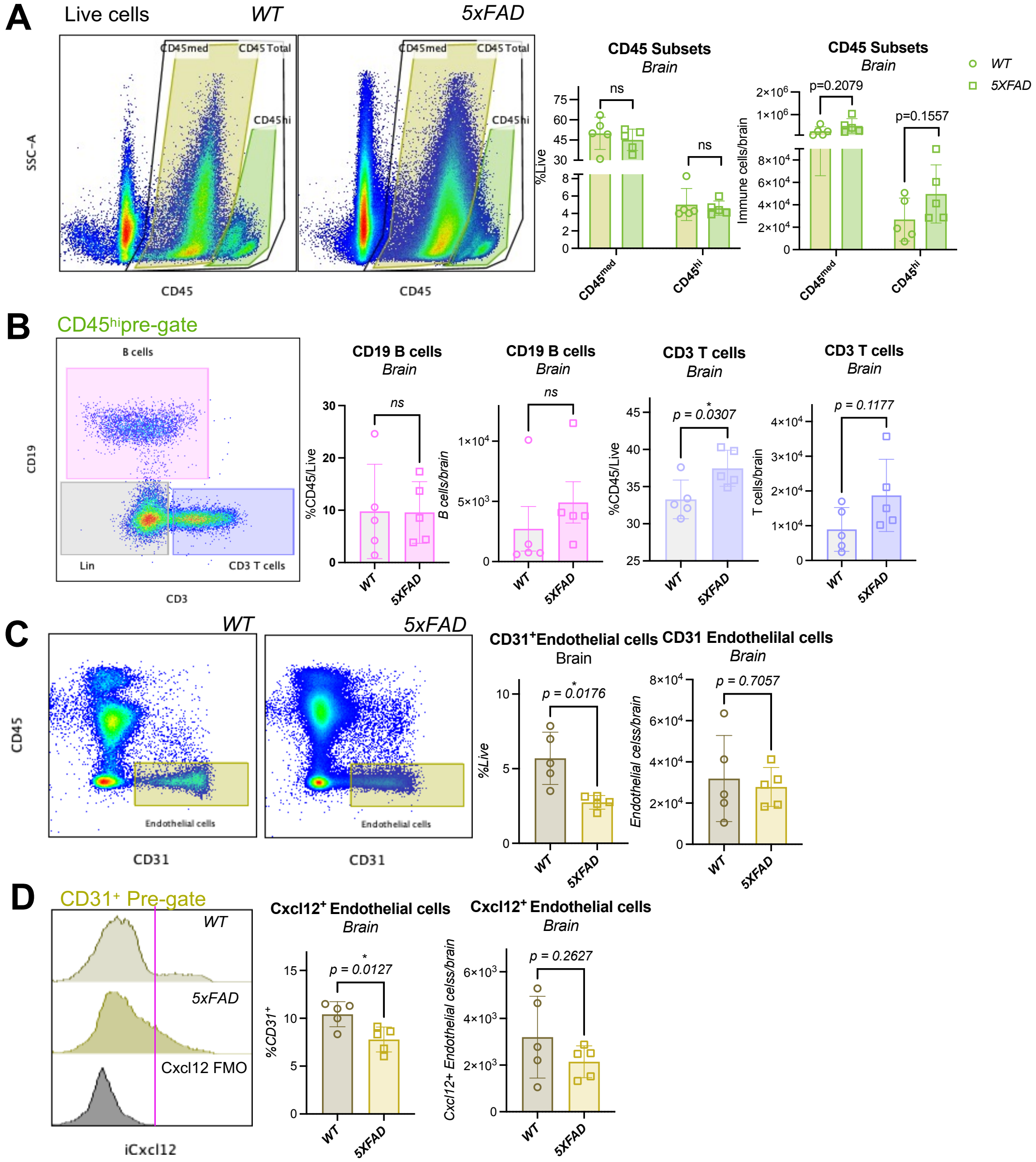

### Fig. S12

**A** *Keren-shaul 2017*

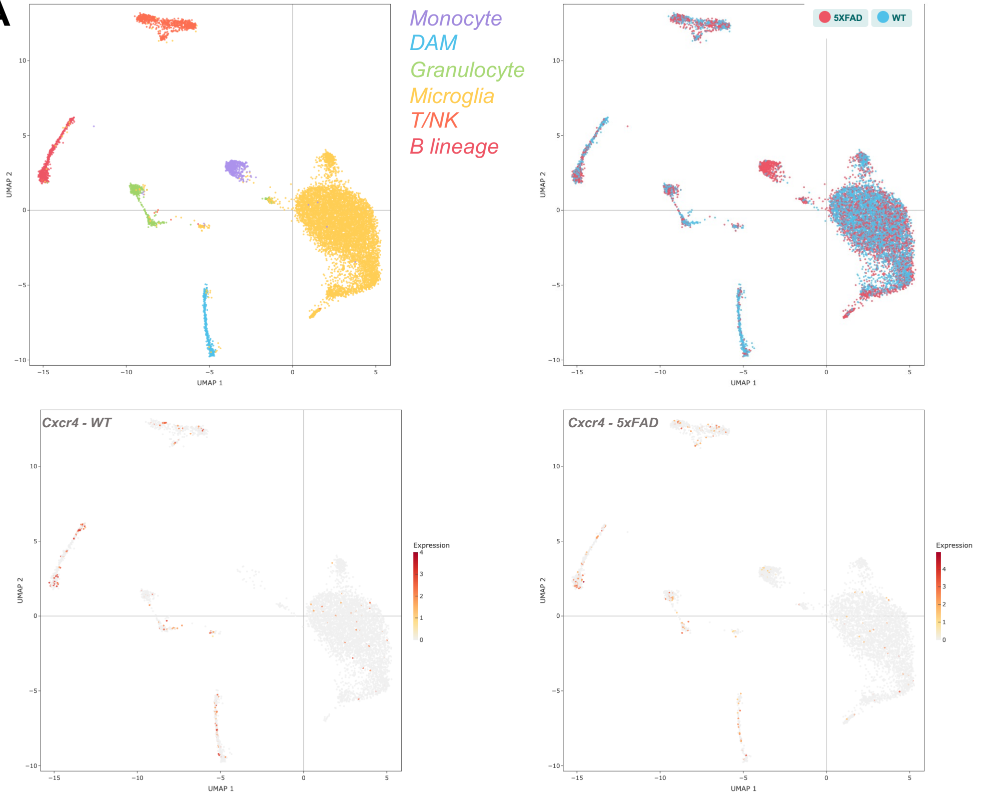


**B** *Su et al 2023*

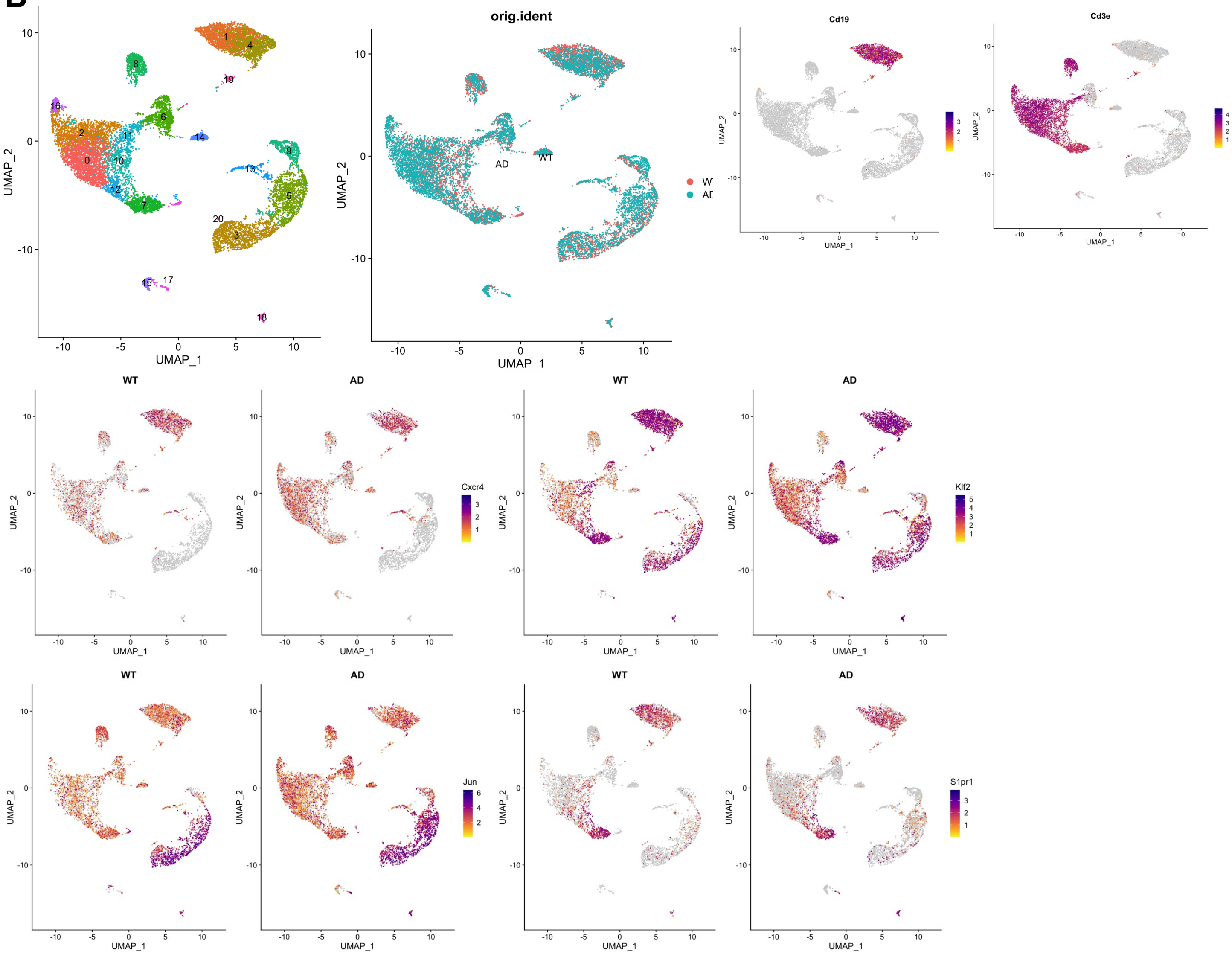

### Fig. S13

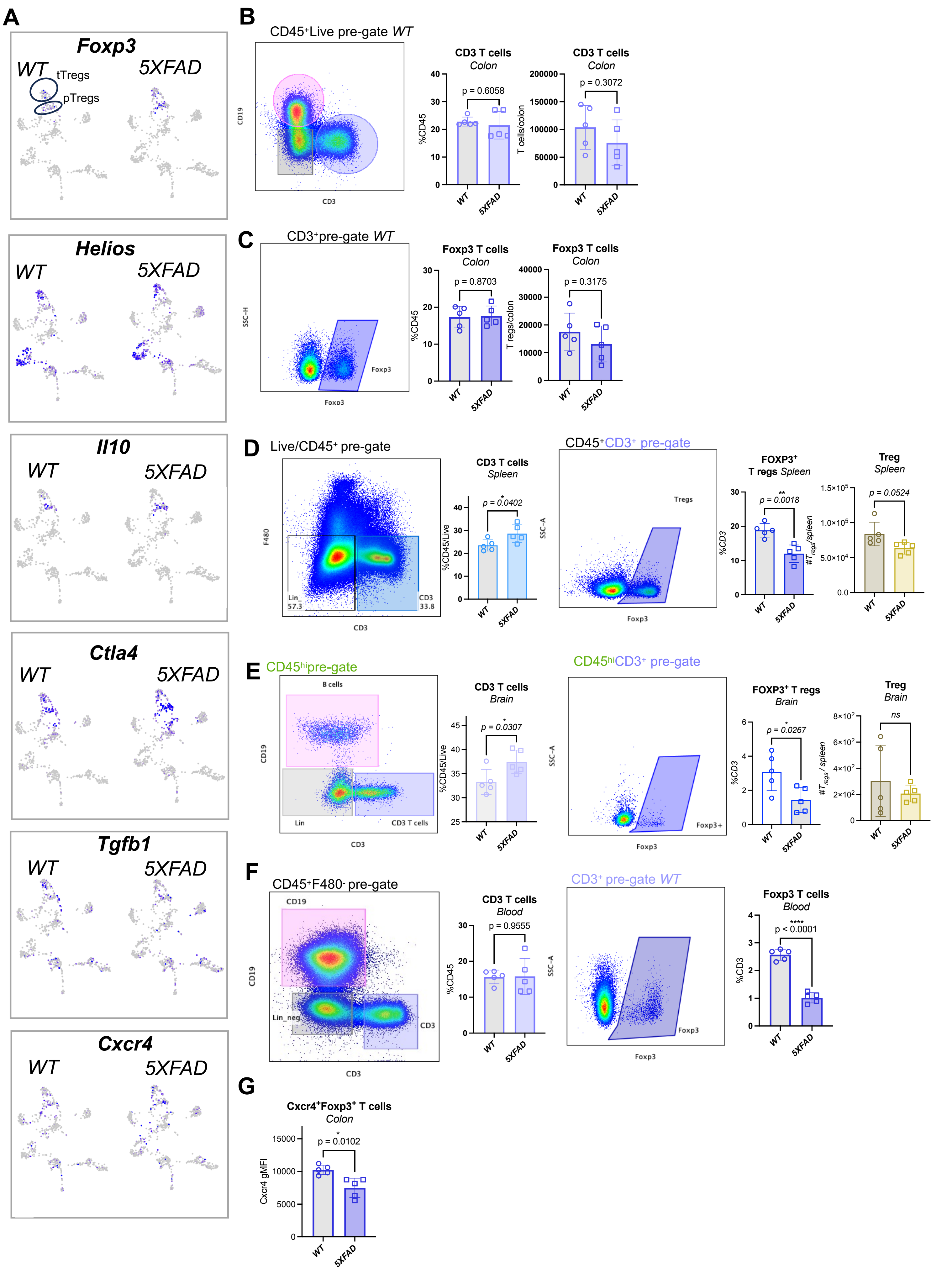

### Fig. S14

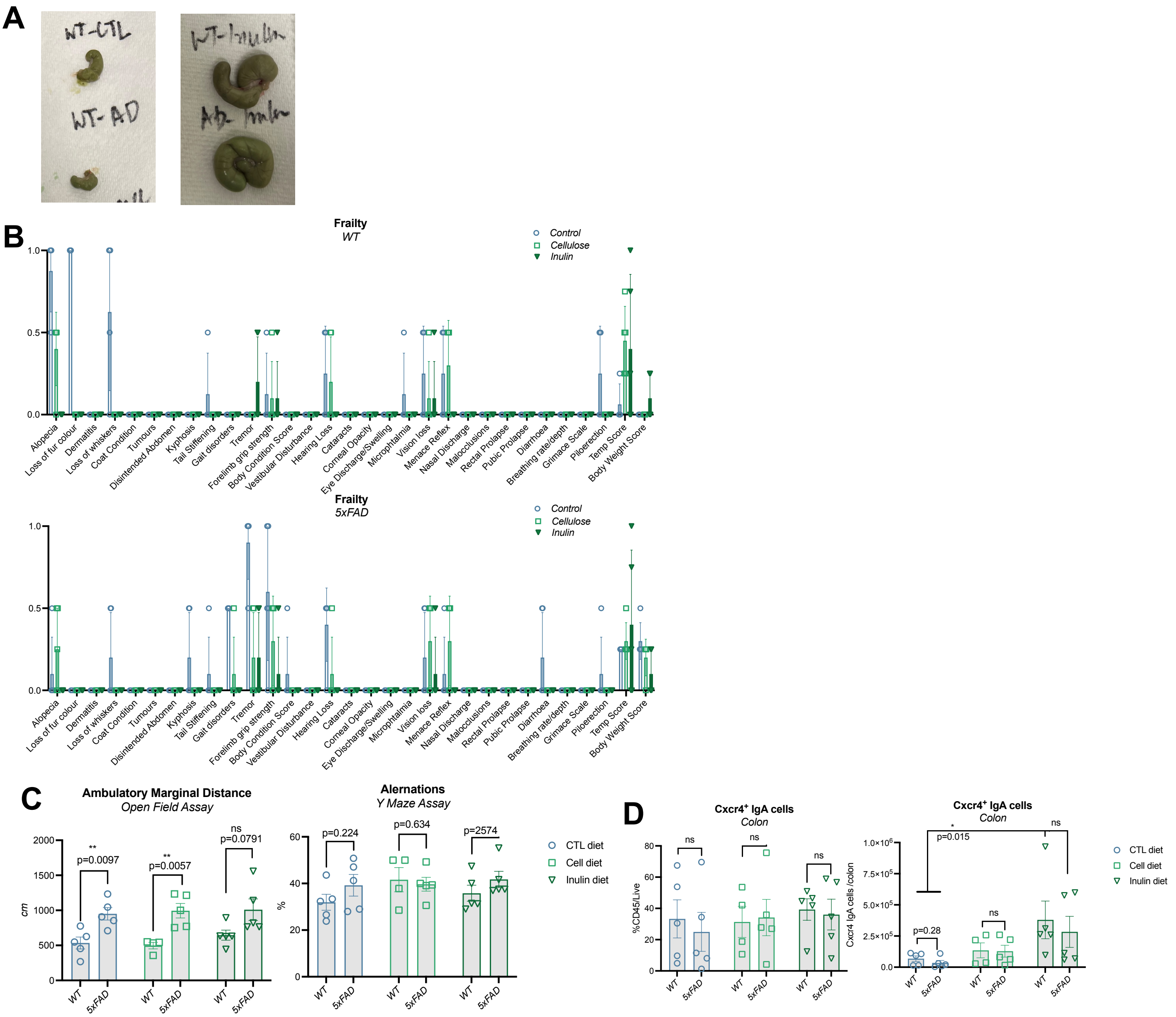
